# Succinate Modulation as a Novel Mechanism Underlying the Effects of Intermittent Fasting on Brain Function and Metabolism in Diet-Induced Obesity

**DOI:** 10.1101/2025.05.16.654006

**Authors:** Andrea Tognozzi, Fabrizia Carli, Sherif Abdelkarim, Sara Cornuti, Francesca Damiani, Maria Grazia Giuliano, Alice Miniati, Martina Nasisi, Lia De Benedictis, Kousha Changizi Ashtiani, Gaia Scabia, Margherita Maffei, Pierre Baldi, Amalia Gastaldelli, Paola Tognini

## Abstract

Obesity significantly impacts the central nervous system (CNS), increasing risks of neuropsychiatric disorders and dementia. Intermittent fasting (IF) shows promise for improving peripheral and CNS health, but its mechanisms are unclear. Using a diet-induced obesity mouse model (10 weeks high fat diet (HFD), then 4 weeks intervention), we compared HFD, HFD-IF, ad libitum control chow (CC), and CC-IF groups. Switching to CC or IF reduced body weight, fat mass, and improved glucose tolerance. Notably, CC-IF uniquely enhanced exploration and reduced anxiety-like behavior. Transcriptomics revealed HFD-induced hippocampal neuroinflammation, while metabolomics identified a specific succinate signature in CC-IF mice: plasma concentration decreased while liver and brown adipose tissue (BAT) levels increased. Succinate supplementation mimicked CC-IF metabolic and behavioral benefits and reduced hippocampal inflammation. These findings suggest that regulating plasma succinate and its metabolism in liver and BAT may represent a novel mechanism underlying the metabolic, neuroinflammatory, and behavioral improvements induced by IF.

## INTRODUCTION

Excessive body weight and obesity are well-established risk factors for the development of numerous comorbidities, including hypertension, type 2 diabetes, stroke, ischemic heart disease, and certain types of cancer.^1–3^ Obesity is also linked to cognitive decline and an increased risk for neurological disorders such as dementia and Alzheimer’s disease (AD).^4–8^ Notably, post-mortem studies of individuals with severe obesity frequently hallmark pathological features of AD, including β-amyloid deposits and β-amyloid precursor proteins.^9^ Furthermore, high body mass index (BMI) has been correlated with deficits in learning, memory, attention, and decision-making.^10–14^ These cognitive impairments are often associated with structural and volumetric changes in the brain, particularly in regions vital for memory and executive function, such as the hippocampus and prefrontal cortex.^15–18^ Preclinical studies consistently demonstrate that western diets, characterized by high-fat and high-sugar intake, as well as high-fat diets (HFD), are associated with a spectrum of cognitive deficits, including impaired learning, memory, increased anxiety- and depression-like behaviours.^19–25^ These effects highlight the detrimental impact of diet-induced obesity (DIO) on brain health, further underscoring the urgent need for effective interventions. While traditional low-calorie balanced diets have long been recognized for their benefits in weight loss and metabolic improvements, recent research suggests that intermittent fasting (IF) may offer distinct and, in some cases, superior advantages. IF, which alternates periods of restricted caloric intake with unrestricted eating, has been shown to improve insulin sensitivity, regulate blood glucose levels, and positively affect lipid profiles in both humans and animal models.^26–33^ In addition to its metabolic benefits fasting extends lifespan,^33,34^ and exerts powerful neuroprotective effects, enhancing cognitive performance, reducing anxiety, and offering protection against depression.^35–40^ Fasting has been found to modulate the transcriptional and epigenetic landscape of the brain, particularly within the cerebral cortex,^41^ suggesting profound, system-wide effects beyond mere metabolic improvements.

Despite the recognized benefits of both dietary approaches, the comparative impact of IF versus low-calorie diets on brain function in the context of obesity remains incompletely understood. IF may activate distinct or complementary molecular pathways that further enhance brain health in a condition of metabolic dysfunction.

In the current study, we aimed to explore how IF and ad libitum consumption of a balanced control chow (CC) diet influence metabolic parameters, cognitive function, and behaviors in a DIO mouse model. Male mice were initially fed a HFD for 10 weeks, followed by random allocation into four experimental groups for an additional 4 weeks: a group continuing on the HFD (HFD group), a group subjected to IF by receiving the HFD every other day (HFD-IF), a group fed a balanced control chow diet ad libitum (CC), and a group subjected to IF with the CC every other day (CC-IF).

Metabolic and behavioural correlates were investigated, revealing that CC-IF was particularly effective in reducing body weight (BW) and improving anxiety-like behaviour. To further elucidate the molecular mechanisms underlying the metabolic and brain related effects of IF, we conducted a transcriptomic experiment on liver and hippocampal tissues. Gene ontology (GO) analysis revealed that inflammatory and immune-related pathways were prominently dysregulated in the hippocampus of HFD mice. Additionally, metabolomic profiling identified a significant increase in succinate levels in the liver and brown adipose tissue (BAT), but a decrease in plasma, specific to CC-IF. To investigate the role of succinate as a potential mediator of CC-IF, we administered sodium succinate to DIO mice and explored its effects on BW reduction, behavior modulation, and hippocampal inflammation.

## RESULTS

### Metabolic effects of dietary switch to intermittent fasting or an ad libitum balanced diet after 10-week HFD feeding

To investigate and compare how dietary interventions, such as IF or a balanced low-calorie diet, affect metabolism in DIO mice, male C57BL/6J mice were fed a HFD for 10 weeks. From week 11 to week 14, mice were subjected to a diet switch (Fig. 1A). BW was measured weekly, and as anticipated, HFD feeding led to a continuous increase in BW from week 1 (Fig. 1B). Transitioning to the HFD-IF, CC, and CC-IF regimens resulted in a significant BW reduction compared to the HFD group (Fig. 1B). Notably, both the CC and CC-IF interventions induced greater weight loss compared to HFD-IF, with the CC-IF group showing a more pronounced reduction during weeks 12 and 13 compared to the CC group (Fig. 1B, see Suppl. Table 1 for the complete statistics).

**Figure 1.**
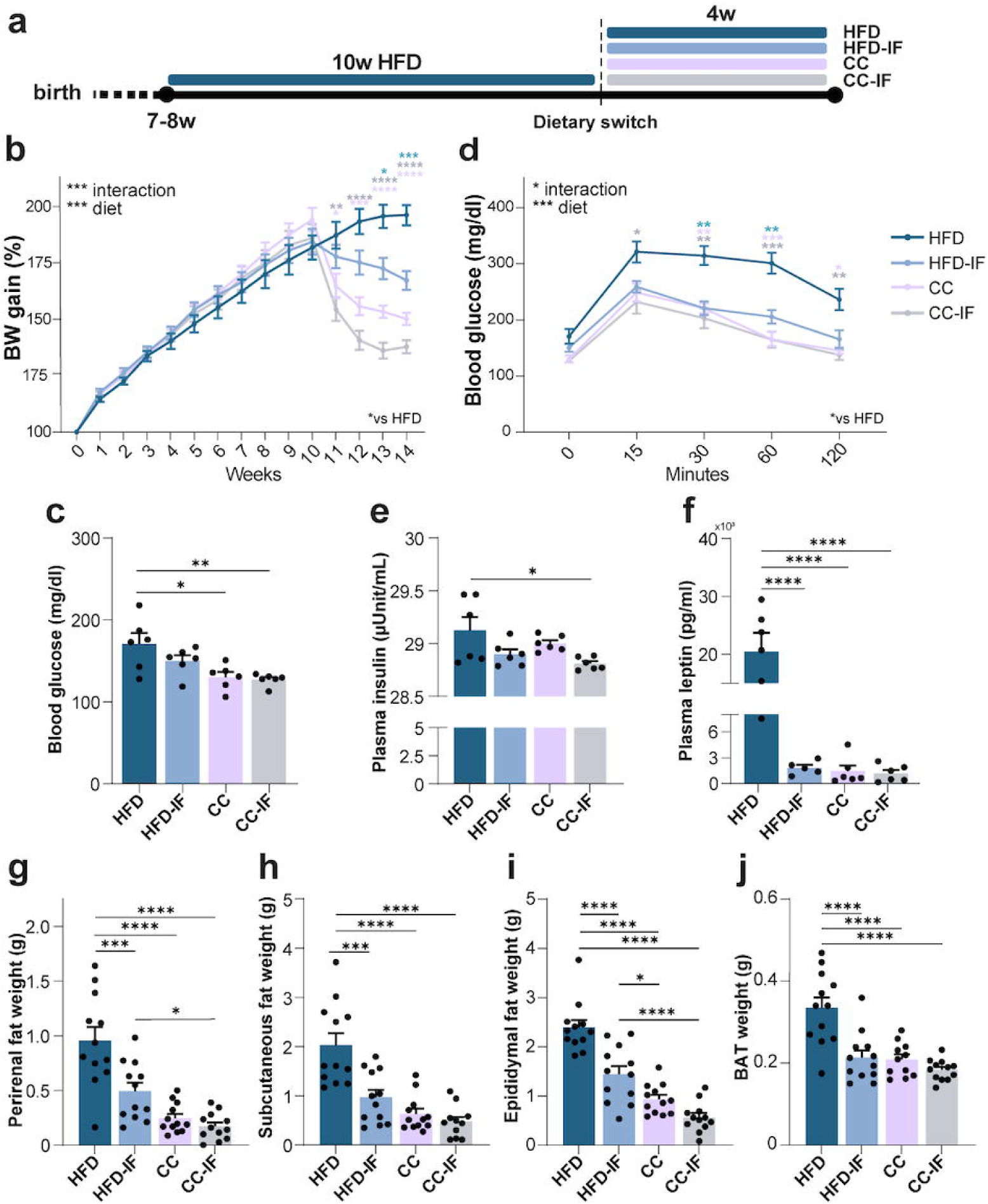
Changes in body weight and metabolic parameters driven by the dietary switch in DIO mice. **A)** Experimental timeline. **B)** Weekly measure of BW (n = 12/group. Two-way RM ANOVA, diet*time interaction p < 0.0001. See Table S1 for Tukey’s post hoc comparisons). **C)** Fasting glycemia (n = 6/experimental group Two-way ANOVA, diet*feeding mode interaction p = 0.2971, feeding mode p = 0.1604, diet p = 0.001. Tukey’s post hoc, HFD vs CC-IF p = 0.0062, HFD vs CC p = 0.0114). **D)** Oral glucose tolerance test (n = 6/experimental group. Two-way RM ANOVA, diet*time interaction p = 0.0206. See Table S1 for Tukey’s post hoc comparisons). **E)** Plasma insulin level (n = 6/experimental group. Two-way ANOVA, diet*feeding mode interaction p = 0.7967, feeding mode p = 0.0073, diet p = 0.1438. Tukey’s post hoc, HFD vs CC-IF p = 0.0221). **F)** Plasma leptin level (n = 5 HFD-IF, n = 6 HFD/CC/CC-IF. Two-way ANOVA, diet*feeding mode interaction p < 0.0001, feeding mode p < 0.0001, diet p < 0.0001. Tukey’s post hoc, HFD vs CC-IF p < 0.0001, HFD vs CC p < 0.0001, HFD vs HFD-IF p < 0.0001). **G)** Perirenal fat weight (n = 12/group. Two-way ANOVA, diet*feeding mode interaction p = 0.0135, feeding mode p = 0.001, diet p < 0.0001. Tukey’s post hoc, HFD vs CC-IF p < 0.0001, HFD vs CC p < 0.0001, HFD vs HFD-IF p = 0.0005, CC-IF vs HFD-IF p = 0.0205). **H)** Subcutaneous fat weight (n = 12/experimental group. Two-way ANOVA, diet*feeding mode interaction p = 0.0081, feeding mode p = 0.0005, diet p < 0.0001. Tukey’s post hoc, HFD vs CC-IF p < 0.0001, HFD vs CC p < 0.0001, HFD vs HFD-IF p = 0.0002). **I)** Epididymal fat weight (n = 12/group. Two-way ANOVA, diet*feeding mode interaction p = 0.0264, feeding mode p < 0.0001, diet p < 0.0001. Tukey’s post hoc, HFD vs CC-IF p < 0.0001, HFD vs CC p < 0.0001, HFD vs HFD-IF p < 0.0001, CC-IF vs HFD-IF p < 0.0001, CC vs HFD-IF p = 0.0332). **J)** Brown adipose tissue weight (n = 12/experimental group. Two-way ANOVA, diet*feeding mode interaction p = 0.0079, feeding mode p = 0.0001, diet p < 0.0001. Tukey’s post hoc, HFD vs CC-IF p < 0.0001, HFD vs CC p < 0.0001, HFD vs HFD-IF p < 0.0001). ANOVA factors: diet = HFD vs CC, feeding mode = ad libitum vs IF. Error bars represent SEM. *p ≤ 0.05, **p ≤ 0.01 and ***p ≤ 0.001, ****p ≤ 0.0001.

During the 14th week, fasting blood glucose levels were measured across all experimental groups. The transition to CC and CC-IF diets resulted in a significant reduction in glycemia compared to HFD-fed mice (Fig. 1C). In contrast, 4 weeks of HFD-IF did not induce a significant decrease in fasting blood glucose (Fig. 1C). To further assess glucose metabolism, we conducted an oral glucose tolerance test (OGTT). After 14 weeks of HFD feeding, mice exhibited glucose intolerance, characterized by elevated blood glucose levels at time 0 and a failure to return to baseline within 2 hours post-glucose administration (Fig. 1D). Notably, the HFD-IF, CC, and CC-IF regimens all effectively restored glucose tolerance following DIO, with no significant differences observed among the three dietary interventions (Fig. 1D, see Suppl. Table 1 for full statistical analysis). Consistent with these findings, HFD-fed mice showed elevated plasma insulin levels, which decreased upon dietary switch and reached significance in CC-IF conditions (Fig. 1E). This suggests that HFD-induced hyperglycemia may be attributed to a loss of insulin sensitivity and subsequent impairment in glucose uptake by specific tissues, potentially driven by increased lipid metabolism.^42,43^ Furthermore, plasma leptin levels were significantly elevated in HFD-fed mice compared to the other three groups (Fig. 1F), due to increased fat depot accumulation.^44^ Specifically, HFD mice exhibited a marked increase in white adipose tissue (WAT) depots, including perirenal, epididymal, and subcutaneous fat, compared to HFD-IF, CC, and CC-IF groups (Fig. 1G-I). Notably, the CC-IF intervention was the most effective in reducing WAT, leading to a significant decrease in both epididymal and perirenal fat relative to the HFD-IF group (Fig. 1G, H). These findings were consistent when evaluating total WAT weight, as well as when normalizing fat depot weights to individual BW (Suppl. Fig. 1). Additionally, interscapular brown adipose tissue (BAT) was significantly reduced to a similar extent across HFD-IF, CC, and CC-IF dietary interventions (Fig. 1J, Suppl. Fig. 1).

All together these data demonstrate that switching to IF or ad libitum CC after 10 weeks HFD feeding had all significant effects on decreasing BW, fat depots and improving glucose tolerance. However, CC-IF seems to be the dietary regimen with the strongest metabolic effects.

### CC-IF ameliorates exploratory and anxiety-like behavior

Neuropsychiatric comorbidities, including anxiety, depression, mood disorders and mild cognitive impairment, are frequent in obese individuals, with a subsequent reduction of the quality of life.^45^ Behavioral abnormalities have also been described in rodent models of DIO. Thus, to gain insights on the potential influence of switching to IF or an ad libitum balanced diet on behavioral performance in comparison to DIO mice, a battery of tests was performed during the last 10 days of feeding (Fig. 2A).

**Figure 2.**
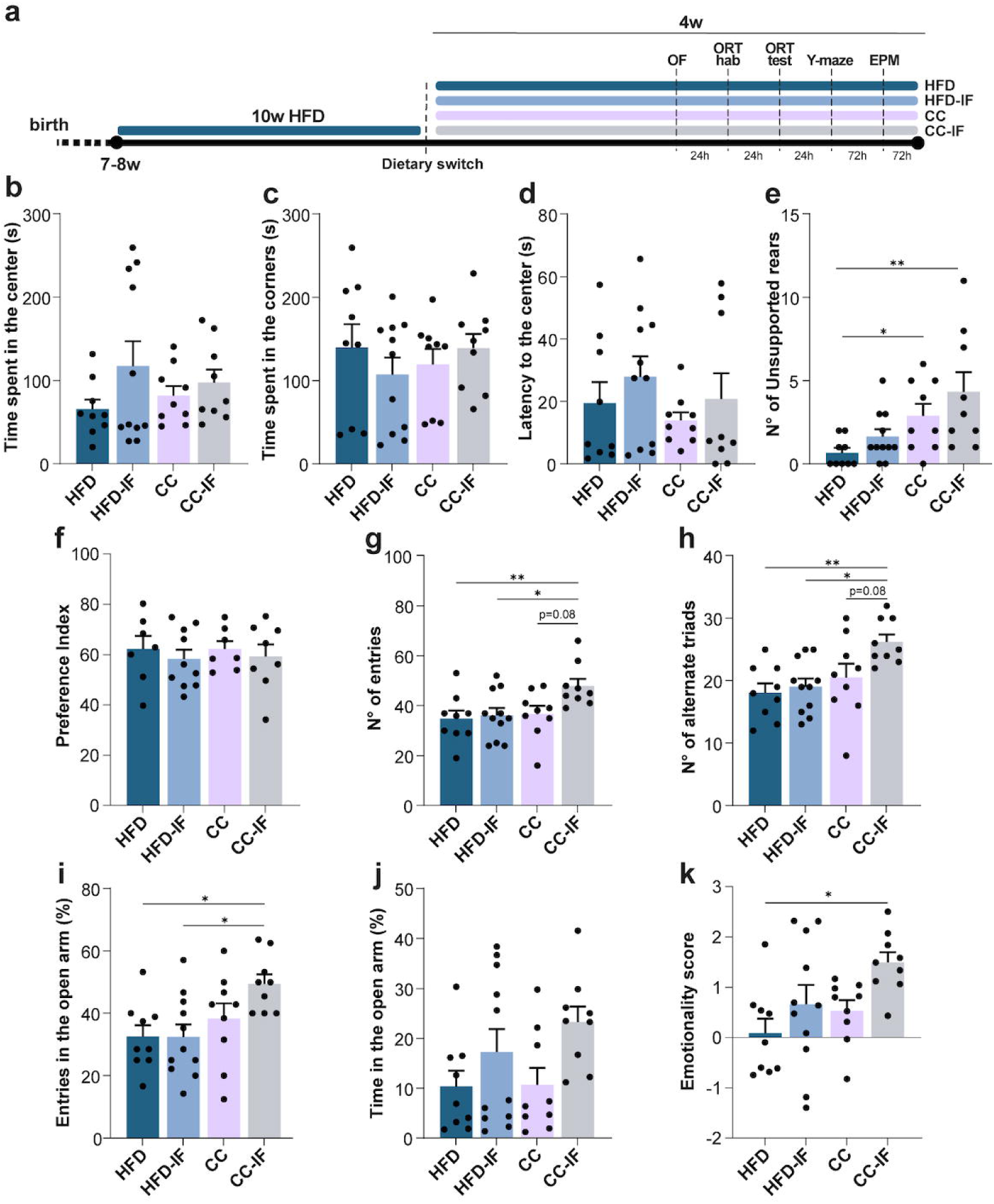
Analysis of behavioural performance after the diet switch in DIO mice. **A)** Experimental timeline. **B)** Time spent in the center in the OF (Two-way ANOVA, diet*feeding mode interaction p = 0.3807, feeding mode p = 0.1076, diet p = 0.922). **C)** Time spent in the corners in the OF (Two-way ANOVA, diet*feeding mode interaction p = 0.2365, feeding mode p = 0.763, diet p = 0.791). **D)** Latency to enter in the central zone in the OF (Two-way ANOVA, diet*feeding mode interaction p = 0.901, feeding mode p = 0.2396, diet p = 0.3348). **E)** Number of unsupported rears (Two-way ANOVA, diet*feeding mode interaction p = 0.7431, feeding mode p = 0.1021, diet p = 0.0016. Tukey’s post hoc, HFD vs CC-IF p = 0.0064, CC-IF vs HFD-IF p = 0.0478). **F)** NORT preference index (Two-way ANOVA, diet*feeding mode interaction p = 0.3355, feeding mode p = 0.7412, diet p = 0.7647). **G)** Number of entries in the Y-maze test (Two-way ANOVA, diet*feeding mode, interaction p = 0.1165, feeding mode p = 0.0494, diet p = 0.0328. Tukey’s post hoc, HFD vs CC-IF p = 0.0282, CC-IF vs HFD-IF p = 0.0418, CC vs CC-IF p = 0.0746. **H)** Y-maze number of alternate triads (Two-way ANOVA, diet*feeding mode, interaction p = 0.1165, feeding mode p = 0.0494, p = 0.0328. Tukey’s post hoc, HFD vs CC-IF p = 0.0058, CC-IF vs HFD-IF p = 0.012, CC vs CC-IF p = 0.0794). **I)** Percentage of entries in the open arm in the EPM test (Two-way ANOVA, diet*feeding mode interaction p = 0.1701, feeding mode p = 0.1758, diet p = 0.0077. Tukey’s post hoc, HFD vs CC-IF p = 0.0306, CC-IF vs HFD-IF p = 0.0209. **J)** EPM percentage of time spent in the open arm (Two-way ANOVA, diet*feeding mode, interaction p = 0.4621, feeding mode p = 0.0150, diet p = 0.405. **K)** Emotionality score. (Two-way ANOVA, diet*feeding mode interaction p = 0.5341, feeding mode p = 0.017, diet p = 0.047. Tukey’s post hoc, HFD vs CC-IF p = 0.0167). N = 8-11/experimental group. Error bars represent SEM. *p ≤ 0.05, **p ≤ 0.01 and ***p ≤ 0.001, ****p ≤ 0.0001.

First, we assessed anxiety-like behavior through the open-field (OF) test. No changes in velocity were observed among the groups (Suppl. Fig. 2A). Moreover, no significant alterations were found in the time spent in the center or in the corner of the arena (Fig. 2B, C) or in the latency to enter in the center (Fig. 2D). Notably, while no differences were present in the total rearing or in the supported rearing (Suppl. Fig. 2B, C), both CC and CC-IF mice performed a higher number of unsupported rearing with respect to the HFD group (Fig. 2E), indicating willingness to explore the new environment and a potential decrease in anxiety.^46^

Declarative memory performance was assessed using the Novel Object Recognition Test (NORT), which leverages the innate tendency of mice to spend more time exploring a novel object compared to a familiar one. After a 24-hour retention interval (testing phase), no significant differences were observed in the preference index among the experimental groups, suggesting that the ability to recognize the familiar object remained intact across all the dietary conditions (Fig. 2F).

The Y-maze test was performed to obtain an estimation of working memory and spontaneous exploratory behavior. The CC-IF group displayed a significant increase in the number of total entries, compared to HFD and HFD-IF, while a trend was identified comparing it to the CC treatment (Fig. 2G). Furthermore, the number of alternate triads of the CC-IF showed the same results (Fig. 2H), suggesting a higher exploratory behavior. On the other hand, no changes were found in the % of alternation parameters, indicating no overt alterations in working memory (Suppl. Fig. 2D).

To further assess exploratory and anxiety-like behaviors, mice were tested in the elevated plus maze (EPM). CC-IF mice exhibited a greater number of entries into the open arm compared to the HFD and HFD-IF groups (Fig. 2I). A main effect of IF was found in the time spent in the open arm (Fig. 2J) compared to the ad libitum groups.

Finally, to provide an overall summary of the behavioral findings, an emotionality score was calculated as an index of anxiety-like behavior.^47^ This score revealed a marked difference between the CC-IF and HFD and HFD-IF mice (Fig. 2K), confirming that CC-IF is the dietary regimen with the stronger effect on anxiety and exploration in comparison to DIO condition.

### Hippocampus transcriptome analysis revealed HFD-driven activation of immune processes and distinct transcriptional responses-driven by the dietary switches

Complex behavior such as exploration and emotional features involves several brain areas, including cortical and subcortical regions. To dissect the mechanisms underlying diet effects on mouse behavioral performance, and based on our results pointing toward an improvement in exploration and anxiety-like behavior in CC-IF, we focused on the hippocampus. RNA-sequencing (seq) experiments of all the four experimental groups revealed significant gene expression differences in the three dietary switches with respect to HFD feeding (Suppl. Table 2). GO analysis (Suppl. Table 2) also highlighted distinct biochemical pathways. A total of 121 transcripts were differentially expressed in HFD vs CC-IF conditions, specifically 28 genes were downregulated and 93 genes were upregulated in HFD (Suppl. Fig. 3A). Among the upregulated transcripts in HFD the top annotations were related to immune system response and processes, antigen processing and presentation, aging (Suppl. Fig. 3B). The genes downregulated in HFD clustered in annotations related to mRNA splicing and translation (Suppl. Fig. 3B). 73 transcripts were altered in the comparison HFD vs CC (Suppl. Fig. 3A), with the 60 genes upregulated in the HFD group again clustering in pathways related to immune system response (Suppl. Fig. 3C). Finally, 135 transcripts were differentially expressed in the hippocampus of HFD vs HFD-IF mice. Antigen processing, response to interferon gamma and aging were again hits in the annotations of the 114 genes upregulated in HFD, while the transcripts upregulated in HFD-IF clustered in mRNA splicing and translation (Suppl. Fig. 3D). To have a better understanding of the common and specific pathways in CC-IF, CC and HFD-IF with respect to HFD feeding, we crossed the differentially expressed transcripts of the 3 conditions in comparison with HFD. The Venn diagram demonstrated that 31 transcripts were commonly dysregulated in the 3 dietary regimens with respect to HFD (Fig. 3B, and Suppl. Table 3), and they clustered in annotations linked to immune system functions (Fig. 3C and Suppl. Table 3). The 60 differentially expressed genes specific to CC-IF clustered in biological pathways linked to protein folding, endoplasmic reticulum stress, and endosomal vesicle fusion (Fig. 3D and Suppl. Table 3). On the other hand the 66 genes specific to HFD-IF response were grouped in annotations such as oxidative phosphorylation, chloride transport and cell growth (Fig. 3E and Suppl. Table 3). CC only genes clustered in pathways involved in metabolism such as glycogen metabolic process, and negative regulation of gluconeogenesis (Fig. 3F and Suppl. Table 3).

**Figure 3.**
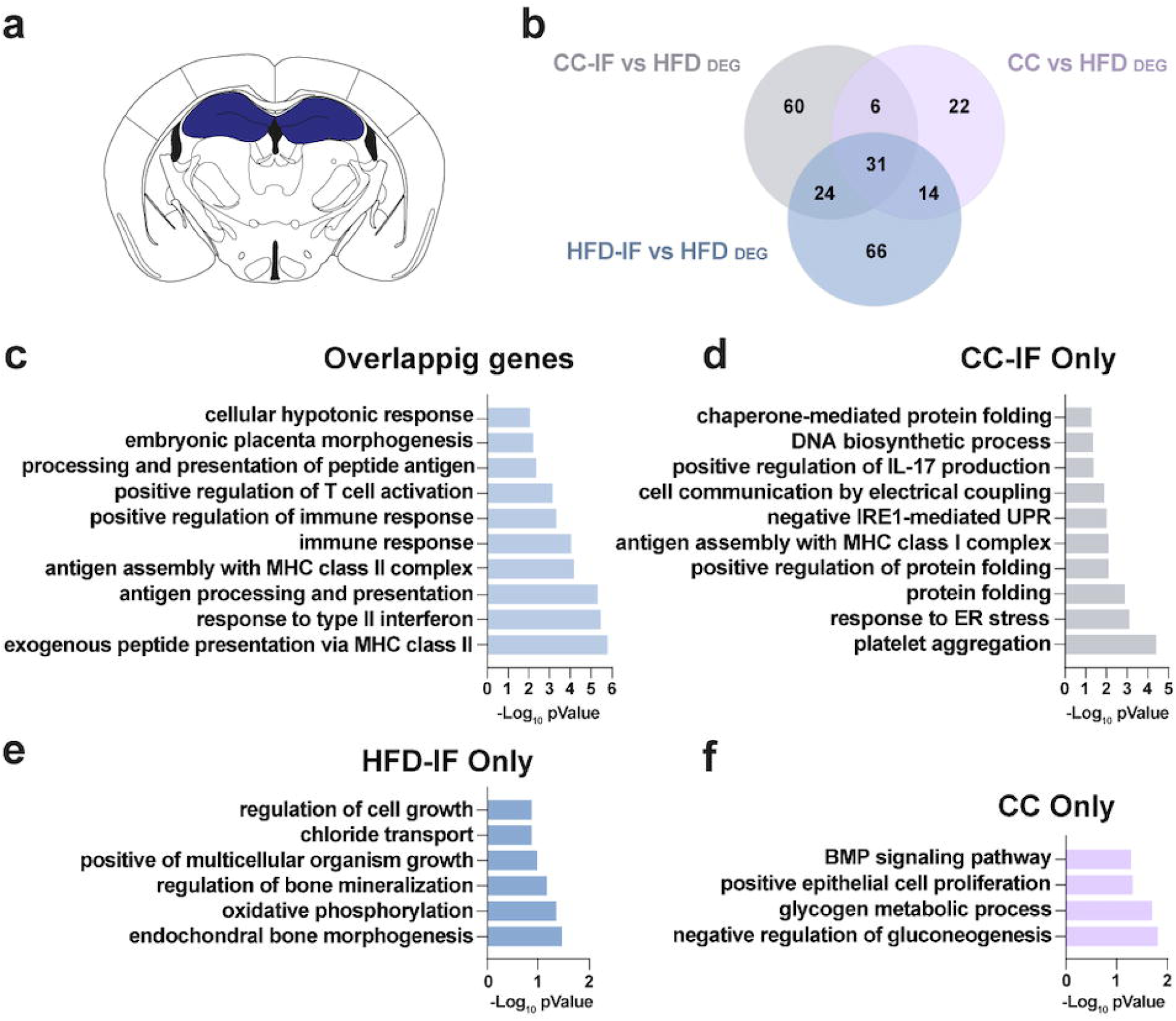
Dietary regimen-dependent transcriptional changes in the hippocampus. **A)** Schematic coronal section of the hippocampus showing the site of RNA-seq analysis. **B)** Venn diagram illustrating the crossing of DEG (pValue < 0.1) between each of the 3 dietary interventions (HFD-IF, CC, CC-IF) compared to HFD. **C)** GO biological processes analysis of 31 overlapping DEG. **D)** GO biological processes analysis of 60 CC-IF exclusive DEG. **E)** GO biological processes analysis of 66 HFD-IF exclusive DEG. **F)** GO biological processes analysis of 22 CC exclusive DEG. n = 4 HFD/HFD-IF/CC, 3 CC-IF.

In summary, the transcriptome analysis revealed that HFD may drive immune response activation and potentially neuroinflammation in the hippocampus of mice. Furthermore, dietary switching to a low calorie diet or IF after 10-week HFD causes hippocampus specific transcriptional reprogramming.

### Liver transcriptome and metabolome analysis revealed specific alterations in TCA cycle metabolites

In parallel to hippocampal RNA-seq, we performed a liver transcriptome analysis (Suppl. Table 4). Liver function is crucial to maintain metabolic homeostasis and it is a key fasting responsive organ, likely mediating many of the beneficial effects of IF.^48,49^ Indeed, hepatic tissue dramatically responded to diet changes. In the comparison HFD vs CC-IF, 768 genes were upregulated in HFD while 769 transcripts were upregulated in CC-IF. In the comparison of HFD vs CC, 274 genes were upregulated in HFD and 189 were upregulated in CC. Finally, HFD liver displayed 281 genes upregulated with respect to HFD-IF, while 296 genes were downregulated (Suppl. Fig. 4A). Importantly, GO analysis (Suppl. Table 4) showed that the transcripts increased by HFD feeding clustered in annotation such as lipid metabolic process, sterol biosynthetic process, steroid biosynthetic and metabolic process, cholesterol metabolic process, independently on the dietary switch (Suppl. Fig. 4B-D). We found common pathways related to circadian rhythms and circadian regulation of gene expression among the transcripts upregulated in the liver of CC-IF, CC and HFD-IF fed mice (Suppl. Fig. 4B-D). As for the hippocampus, we crossed the differentially expressed transcripts of the 3 dietary switches (CC, CC-IF, and HFD-IF) in comparison with HFD (Fig. 4A). The top annotation of the 81 common genes belonged to pathways linked to fatty acid, cholesterol metabolism. Response to glucose, circadian rhythms and insulin secretion also came up in the analysis (Fig. 4B and Suppl. Table 5). The CC-IF vs HFD 1066 specific transcripts displayed hits related to regulation of transcription, translation and chromatin organization (Fig. 4C and Suppl. Table 5). The 242 differentially expressed genes specific to the comparison HFD vs CC clustered in fatty acid and glutathione metabolic processes, response to stress (Fig. 4D and Suppl. Table 5). The 202 transcripts specific to the comparison HFD-IF vs HFD were related to extracellular matrix organization, cell migration, cell adhesion and apoptosis (Fig. 4E and Suppl. Table 5).

**Figure 4.**
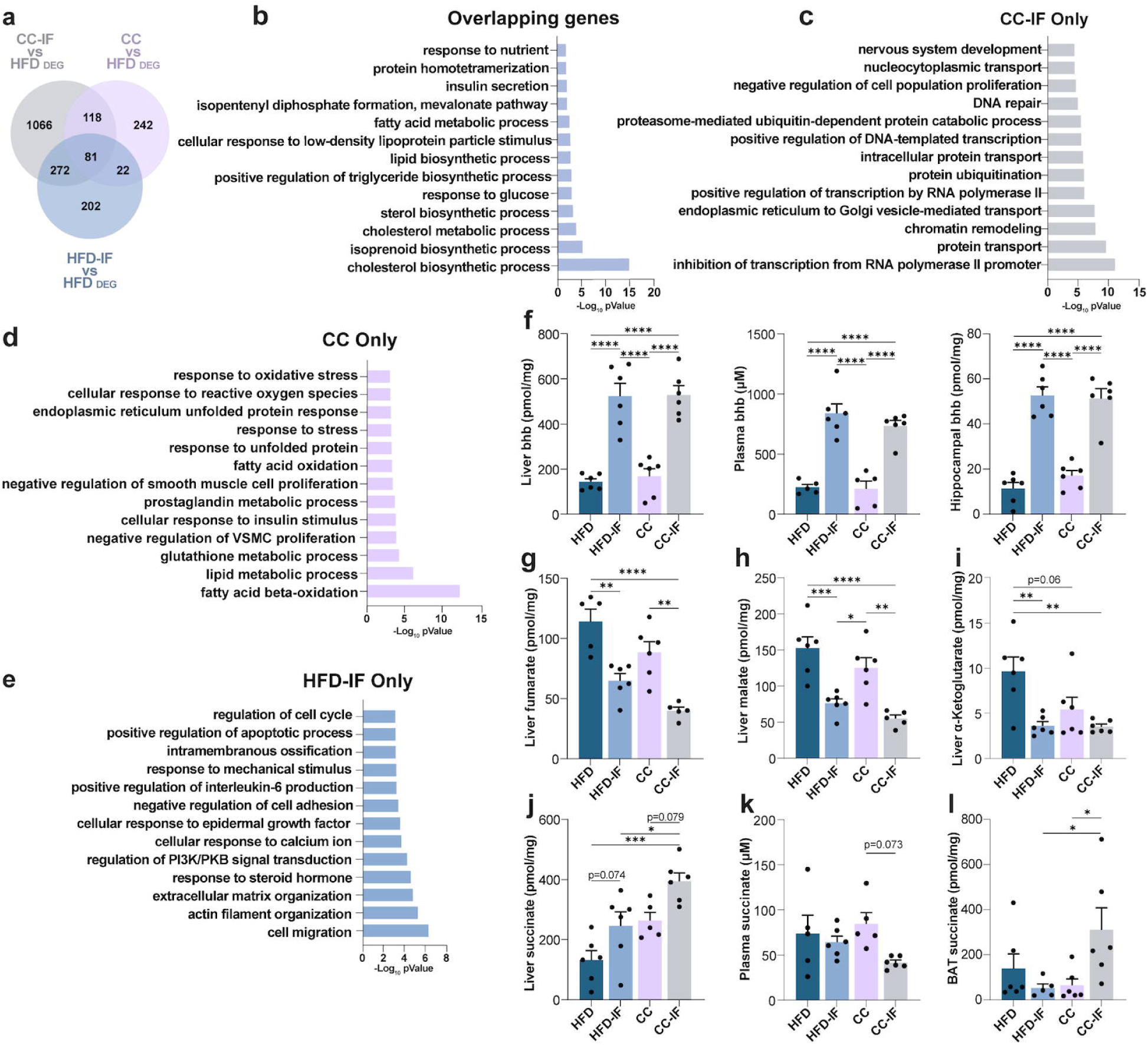
Dietary regimen-dependent transcriptional changes in the liver and metabolomics analysis. **A)** Venn diagram illustrating the crossing of DEG (pValue < 0.05) between each of the 3 dietary interventions (HFD-IF, CC, CC-IF) in comparison with HFD. **B)** GO biological processes analysis of 81 overlapping DEG. **C)** GO biological processes analysis of 1066 CC-IF exclusive DEGs. **D)** GO biological processes analysis of 242 CC exclusive DEG. **E)** GO biological processes analysis of 202 HFD-IF exclusive DEG (n = 4 HFD/CC/CC-IF, n = 2 HFD-IF). **F)** β-hydroxybutyrate concentration in liver, plasma and hippocampus. Liver: Two-way ANOVA, diet*feeding mode interaction p = 0.812, feeding mode p < 0.0001, diet p = 0.7016. Tukey’s post hoc, HFD vs CC-IF p < 0.0001, HFD, HFD vs HFD-IF p < 0.0001, CC vs HFD-IF p < 0.0001, CC vs CC-IF p < 0.0001; Plasma: Two-way ANOVA, diet*feeding mode interaction p = 0.4413, feeding mode p < 0.0001, diet p = 0.3307. Tukey’s post hoc, HFD vs CC-IF p < 0.0001, HFD vs HFD-IF p < 0.0001, CC vs HFD-IF p < 0.0001, CC vs CC-IF p < 0.0001; Hippocampus: Two-way ANOVA, diet*feeding mode interaction p = 0.3063, feeding mode p < 0.0001, diet p = 0.5214. Tukey’s post hoc, HFD vs CC-IF p < 0.0001, HFD vs HFD-IF p < 0.0001, CC vs HFD-IF p < 0.0001, CC vs CC-IF p < 0.0001. **G)** Liver fumarate concentration (Two-way ANOVA, diet*treatment interaction p = 0.9755, feeding mode p < 0.0001, diet p = 0.004. Tukey’s post hoc, HFD vs CC-IF p < 0.0001, HFD vs CC p = 0.1223, HFD vs HFD-IF p = 0.0014, CC vs CC-IF p = 0.0015). **H)** Liver Malate concentration (Two-way ANOVA, diet*feeding mode interaction p = 0.7982, feeding mode p < 0.0001, diet p = 0.0524. Tukey’s post hoc, HFD vs CC-IF p < 0.0001, HFD vs HFD-IF p = 0.008, CC vs HFD-IF p = 0.0313, CC vs CC-IF p = 0.0028). **I)** Liver α-Ketoglutarate concentration (Two-way ANOVA, diet*feeding mode interaction p = 0.0741, feeding mode p = 0.0018, diet p = 0.0624. Tukey’s post hoc, HFD vs CC-IF p = 0.0041, HFD vs CC p = 0.0578, HFD vs HFD-IF p = 0.0047). **J)** Liver Succinate concentration (Two-way ANOVA, diet*feeding mode interaction p = 0.8101, feeding mode p = 0.0024, diet p = 0.0008. Tukey’s post hoc, HFD vs CC-IF p = 0.0002, HFD vs CC p = 0.0764, CC-IF vs HFD-IF p = 0.029, CC vs CC-IF p = 0.0788). **K)** Plasma succinate concentration (Two-way ANOVA, diet*feeding mode interaction p = 0.1638, feeding mode p = 0.0357, diet p = 0.6147. Tukey’s post hoc, CC vs CC-IF p = 0.0731). **L)** BAT succinate concentration (Two-way ANOVA, diet*feeding mode interaction p = 0.017, feeding mode p = 0.2255, diet p = 0.1651. Tukey’s post hoc, CC-IF vs HFD-IF p = 0.0507, CC vs CC-IF p = 0.0508. Other comparisons were not significant. N= 5-6/experimental group. Error bars represent SEM. *p ≤ 0.05, **p ≤ 0.01 and ***p ≤ 0.001, ****p ≤ 0.0001.

Finally, we performed a metabolome analysis of liver, hippocampus and plasma samples focusing on the TCA cycle metabolites and beta-hydroxy-butyrate (bhb). Not surprisingly, bhb was significantly increased in CC-IF and HFD-IF with respect to HFD and CC in liver, plasma and hippocampus (Fig. 4F). Fumarate, malate and alpha-keto-glutarate were significantly decreased in both CC-IF and HFD-IF liver in comparison to HFD (Fig. 4G-I). Those results suggest an increase in TCA cycle activity in HFD liver, suggesting a potential increase in oxidative metabolism and subsequent oxidative stress.^50^ Pyruvate and lactate displayed the same trend (Suppl. Fig. 4E), suggesting an effect of fasting on the enhancement of the gluconeogenic pathway independently on the type of food consumed. Moreover CC-IF fumarate and malate were higher compared to CC (Fig. 4G, H). Interestingly, succinate was the only metabolite among those analyzed to show an exclusive increase in the liver under the CC-IF condition (Fig. 4J), and its levels were inversely correlated to BW (Suppl. Fig. 4F). This observation prompted us to extend the analysis of TCA cycle metabolites to plasma and brain tissues (Suppl. Fig. 4G, H). In plasma samples malate and alpha-keto-glutarate were significantly increased in CC compared to HFD (Suppl. Fig. 4G). No differences were observed in pyruvate due to high variability, while plasma lactate reflected the liver situation (Suppl. Fig. 4G). In the hippocampus, fumarate, malate and alpha-keto-glutarate displayed the same trend with an increase induced by the three dietary switches compared to HFD, although only malate reached statistical significance (Suppl. Fig. 4H). Pyruvate was unchanged, while lactate increased in CC with respect to CC-IF and HFD (Suppl. Fig. 4H). Hippocampal succinate was not altered (Suppl. Fig. 4H). Intriguingly, succinate levels in plasma were affected, showing a decrease in CC-IF (Fig. 4K). Additionally, BAT succinate displayed a distinct increase in CC-IF mice (Fig. 4L). These findings suggest a potential CC-IF-driven “sink effect” for circulating succinate, which may be utilized for thermogenesis in BAT,^51^ and possibly to provide the energy required for gluconeogenesis in the liver.

### Succinate administration improved metabolic parameters in DIO mice

Succinate is a versatile molecule acting as a pivotal constituent in metabolic pathways and as a signaling metabolite in a paracrine and endocrine manner influencing cell responses to a variety of stimuli.^52^ Since the increase in liver and BAT succinate, and its decrease in plasma seemed to be a signature of weight loss and CC-IF, we decided to further explore the succinate potential effects on mouse metabolism and behavioral performance in our DIO model. To this end, after 10 weeks of HFD feeding, succinate dissolved in drinking water was administered to HFD and CC fed mice for the subsequent 4 weeks (Fig. 5A). BW gain was monitored throughout the 14-week experiment. During the first 10 weeks of ad libitum HFD, all groups showed a steady increase in BW (Fig. 5B). From week 11 to week 14, the HFD group continued to gain weight while the CC group showed a marked reduction in BW. The CC-Succ group also displayed a significant BW decrease with respect to the previous HFD condition and in comparison to both HFD and HFD-Succ groups (Fig. 5B). Notably, CC-Succ mice displayed a stronger drop in the BW with respect to CC, similarly to what we observed with the CC-IF intervention (Fig. 1B), and the further decrease in BW was significant with respect to CC by week 14 (Fig. 5B). Finally, the HFD-Succ group showed a slight reduction in BW compared to the HFD group, suggesting that succinate supplementation may have an effect in mitigating the weight gain associated with a HFD (Fig. 5B, for the complete statistics see Suppl. Table 6).

**Figure 5.**
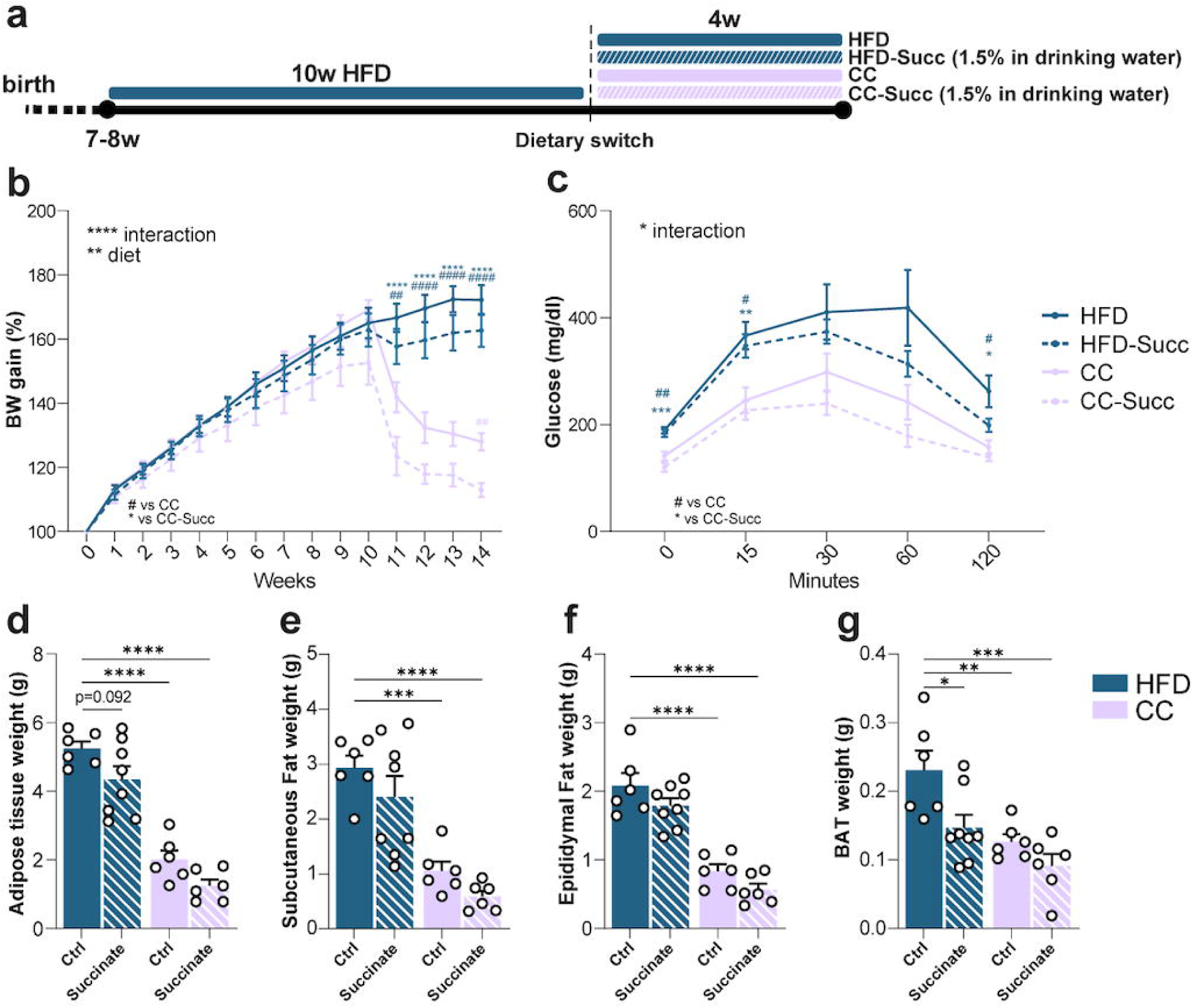
Succinate administration affects body weight, glucose tolerance and fat depots. **A)** Experimental timeline. Mice were fed HFD for 10 weeks and then divided into four experimental groups for the following 4 weeks: one group continued on the HFD drinking regular water (HFD), another group was fed HFD drinking sodium succinate dissolved in water (HFD-Succ), a third group switched to CC drinking regular water (CC), and the fourth group was fed CC drinking sodium succinate (CC-Succ). **B)** Weekly BW measure (n = 11 HFD, n = 14 HFD-Succ, n = 11 CC, n = 12 CC-Succ. Two-way RM ANOVA, diet*time interaction p < 0.0001. See Table S2 for Tukey’s post hoc comparisons). **C)** Oral glucose tolerance test (Two-way RM ANOVA, diet*time interaction p = 0.0206. See Table S2 for Dunnett’s post hoc comparisons). **D)** Total adipose tissue weight (Two-way ANOVA, diet*treatment interaction p = 0.8379, treatment p = 0.01, diet p < 0.0001. Dunnett’s post hoc, HFD vs HFD-Succ p = 0.0921, HFD vs CC < 0.0001, HFD vs CC-Succ p < 0.0001). **E)** Subcutaneous fat weight (Two-way ANOVA, diet*treatment interaction p = 0.9129, treatment p = 0.0822, diet p < 0.0001. Dunnett’s post hoc, HFD vs HFD-Succ p = 0.0921, HFD vs CC p < 0.0001, HFD vs CC-Succ p < 0.0001). **F)** Epididymal fat weight (Two-way ANOVA, diet*treatment interaction p = 0.9573, treatment p = 0.039, diet p < 0.0001. Dunnett’s post hoc, HFD vs HFD-Succ p = 0.2573, HFD vs CC p < 0.0001, HFD vs CC-Succ p < 0.0001). **F)** BAT (Two-way ANOVA, diet*treatment interaction p = 0.2416, treatment p = 0.039, diet p = 0.0007. Dunnett’s post hoc, HFD vs HFD-Succ p = 0.0156, HFD vs CC p = 0.0051, HFD vs CC-Succ p = 0.0003). The label Control (Ctrl) under the graph bars refers to mice drinking regular water. ANOVA factors: diet = HFD vs CC, treatment = succinate vs regular water. N = 5-8/experimental group, unless otherwise stated. Error bars represent SEM. *p ≤ 0.05, **p ≤ 0.01 and ***p ≤ 0.001, ****p ≤ 0.0001.

Glucose tolerance was impaired in HFD. Glycemia decreased 2 hours post-glucose administration in HFD, although it remained significantly higher with respect to CC and CC-Succ (Fig. 5C, for the complete statistics see Suppl. Table 6). In contrast, CC and CC-Succ had lower fasting blood glucose levels, and at 15 minutes post glucose administration in comparison with the HFD groups (Fig. 5C), indicating an improved glucose tolerance. The HFD-Succ curve had a different trend in comparison to HFD. Indeed glycemia started to decrease 30 minutes after glucose administration, a shared characteristic with the CC groups curve, and went back to the fasting levels after 120 minutes (Fig. 5C). This data suggest that succinate supplementation could ameliorate glucose tolerance in HFD fed mice. Importantly, the CC-Succ group consistently showed the most efficient glucose clearance throughout the test, returning to fasting levels by 60 minutes post-administration (Fig. 5C).

Fat depot weight demonstrated the drastic effect of switching to CC. Indeed, total fat, subcutaneous fat, epididymal fat and BAT were significantly decreased in both CC groups in comparison to HFD groups (Fig. 5D-G and see Suppl. Fig. 5A for the analysis after normalization to the BW). Notably, there was a certain effect of succinate supplementation on total fat weight, showing a decrease (Fig. 5D). Subcutaneous and epididymal fat displayed a slight trend toward a decrease (Fig. 5E-F), while BAT was particularly influenced by succinate supplementation in the HFD comparison but not in the CC comparison (Fig. 5G).

### Succinate supplementation improved exploration and anxiety-like behavior similarly to CC-IF

To investigate the potential effect of succinate administration on brain function and evaluate similarities to CC-IF regimen, mice were subjected to behavioral tests during the 14th week of diet (Fig. 6A).

**Figure 6.**
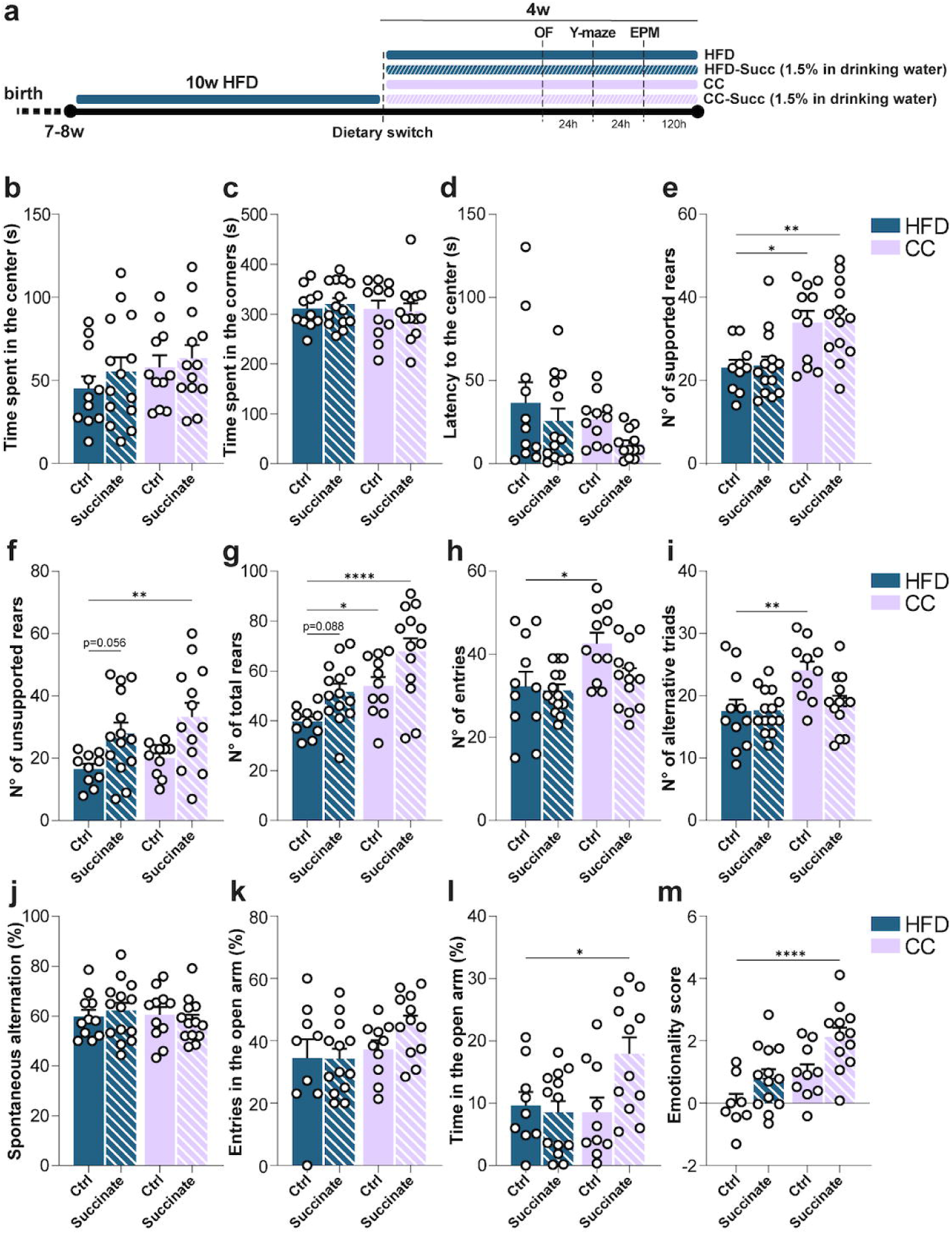
Succinate administration improves behavioral parameters. **A)** Experimental timeline. **B)** Time spent in the center in the OF (Two-way ANOVA, diet*treatment interaction p = 0.7756, treatment p = 0.3318, diet p = 0.1994). **C)** Time spent in the corners in the OF (Two-way ANOVA, diet*treatment interaction p = 0.6295, treatment p = 0.8741, diet p = 0.5938. **D)** Latency to enter in the central zone in the OF (Two-way ANOVA, diet*treatment interaction p = 0.7854, treatment p = 0.0947, diet p = 0.1161). **E)** Number of supported rears (Two-way ANOVA, diet*treatment interaction p = 0.9858, treatment p = 0.8097, diet p < 0.0001. Dunnett’s post hoc, HFD vs HFD-Succ p = 0.9969, HFD vs CC p = 0.0126, HFD vs CC-Succ p = 0.0057). **F)** Number of unsupported rears (Two-way ANOVA, diet*treatment interaction p = 0.7895, treatment p = 0.0007, diet p = 0.2002. Dunnett’s post hoc, HFD vs HFD-Succ p = 0.056, HFD vs CC p = 0.8215, HFD vs CC-Succ p = 0.0038. **G)** Number of total rears (Two-way ANOVA, diet*treatment interaction p = 0.8076, treatment p = 0.0018, diet p = 0.0003. Dunnett’s post hoc, HFD vs HFD-Succ p = 0.0883, HFD vs CC p = 0.0462, HFD vs HFD-Succ p < 0.0001. **H)** Y-maze number of entries in the arms. (Two-way ANOVA, diet*treatment interaction p = 0.1550, treatment p = 0.0704, diet p = 0.0078. Dunnett’s post hoc, HFD vs HFD-Succ p = 0.9825, HFD vs CC p = 0.0176, HFD vs CC-Succ p = 0.0176). **I)** Y-maze number of alternate triads (Two-way ANOVA, diet*treatment, interaction p = 0.0514, treatment p = 0.06, diet p = 0.0069. Dunnett’s post hoc, HFD vs HFD-Succ p > 0.9999, HFD vs CC p = 0.0059, HFD vs CC-Succ p = 0.8568). **J)** Y-maze percentage of spontaneous alternation (Two-way ANOVA, diet*treatment interaction p = 0.4118, treatment p = 0.9954, diet p = 0.5732). **K)** EPM percentage of entries in the open arm (Two-way ANOVA, diet*treatment, interaction p = 0.2826, treatment p = 0.3106, diet p = 0.0764). **L)** EPM percentage of time spent in the open arm (Two-way ANOVA, diet*treatment interaction p = 0.0293, treatment p = 0.0809, diet p = 0.0806. Dunnett’s post hoc, HFD vs HFD-Succ p = 0.9736, HFD vs CC p = 0.9778, HFD vs CC-Succ p = 0.0447. **M)** Emotionality score (Two-way ANOVA, diet*treatment interaction p = 0.5805, treatment p = 0.002, diet p = 0.0004. Dunnett’s post hoc, HFD vs HFD-Succ p = 0.1491, HFD vs CC p = 0.088, HFD vs CC-Succ p < 0.0001). N = 11-14/experimental group. Error bars represent SEM. *p ≤ 0.05, **p ≤ 0.01 and ***p ≤ 0.001, ****p ≤ 0.0001.

In the OF test, there was a HFD effect on velocity, demonstrating a reduction (Suppl. Fig. 5B). There were no significant changes in the time spent in the center, in the corner of the arena (Fig. 6B, C) or in the latency to enter in the center (Fig. 6D). Supported rearing was increased by CC with respect to HFD, with no succinate influence (Fig. 6E). When analyzing unsupported rearing behavior, we observed a notable effect of succinate supplementation which increased the number of rears in both CC and HFD-fed mice (Fig. 6F). Finally, the total number of rears were increased in CC-Succ and CC compared to HFD, while a tendency to perform a higher number was present in HFD-Succ (Fig. 6G). This data suggests that succinate may improve exploration and reduce overall anxiety-like behavior, regardless of the diet consumed by the animals.

In the Y-maze test, velocity reflected the previous observations from the OF test (Suppl. Fig. 5C). We observed a significant effect of the CC diet switch on the number of entries performed by the mice (Fig. 6H). Succinate supplementation seemed to decrease the number of alternative triads performed by CC mice (Fig. 6I), however the percentage of alternation was affected neither by diet nor succinate (Fig. 6J).

In the EPM, there were no differences in the four groups’ velocity of exploration (Suppl. Fig. 5D). Neither diet nor succinate had effects on the number of entries into the open arm (Fig. 6K). However, succinate supplementation to CC mice increased the time spent in the open arm compared to the HFD group (Fig. 6L), suggesting a reduction in anxiety. Finally, CC-Succ displayed the higher emotionality score (Fig. 6M), confirming the single behavioural test results. Overall, the behavioral data suggest that succinate may reduce anxiety-like behavior and improve exploration, particularly when transitioning to a CC diet, similar to the effects of IF on a CC diet.

### Succinate supplementation affects inflammatory marker levels in the hippocampus

Since the transcriptome analysis revealed that HFD may drive immune response activation and potentially neuroinflammation in the mouse hippocampus, we explored the expression of specific inflammatory markers. Hippocampal IL-1β expression analysis revealed a significant reduction with succinate supplementation, particularly when comparing CC to CC-Succ (Fig. 7A). Similarly, TNFα expression was significantly lower in mice switched to a CC diet with succinate treatment, and a reduction trend was observed in HFD-Succ mice (Fig. 7B).

**Figure 7.**
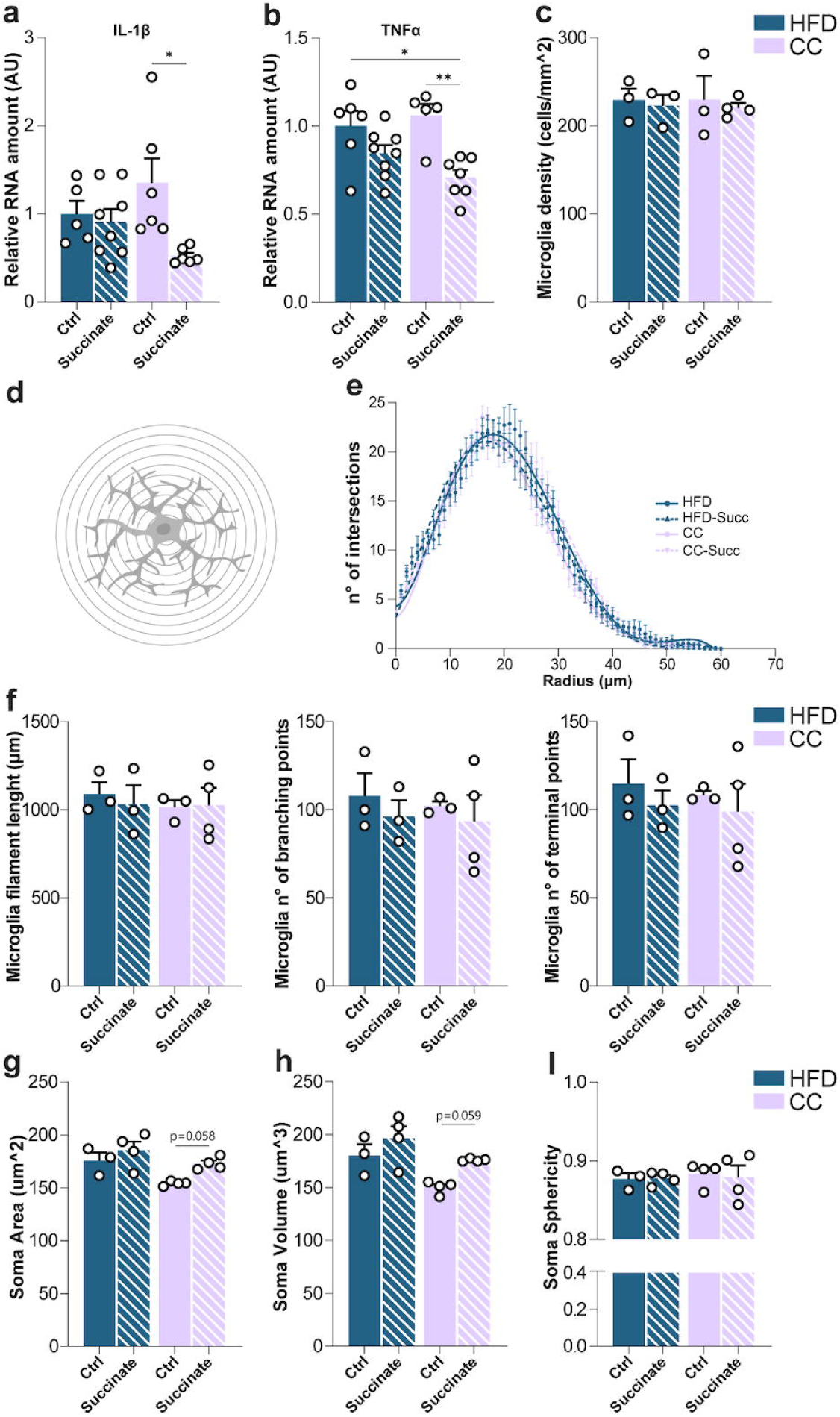
Succinate administration affects hippocampal neuroinflammatory markers. **A**) IL-1ꞵ Relative mRNA expression (Two-way ANOVA, diet*treatment interaction p = 0.0473, treatment p = 0.0168, diet p = 0.9293. Tukey’s post hoc, CC vs CC-Succ p = 0.0163). **B)** TNF⍺ Relative mRNA expression (Two-way ANOVA, diet*treatment interaction p = 0.124, treatment p = 0.0004, diet p = 0.5353. Tukey’s post hoc, HFD vs CC-Succ p = 0.0124, CC vs CC-Succ p = 0.004). **C)** Density of hippocampal microglia cells (Two-way ANOVA, diet*treatment interaction p = 0.9287, treatment p = 0.6269, diet p = 0.9455. **D)** Representative image of a Sholl analysis for hippocampal microglia reconstruction. **E)** Sholl analysis (n=14/20 cells per group. Two-way ANOVA, distance*treatment interaction p = 0.4595, distance p < 0.0001, treatment p = 0.5802). **F)** Microglia Filament length (Two-way ANOVA, diet*treatment interaction p = 0.6954, treatment p = 0.795, diet p = 0.6602). N° of branching points (Two-way ANOVA, diet*treatment interaction p = 0.9032, treatment p = 0.4183, diet p = 0.7261). N° of terminal points (Two-way ANOVA, diet*treatment interaction p = 0.9095, treatment p = 0.4107, diet p = 0.6927). **G)** Microglia soma area (Two-way ANOVA, diet*treatment interaction p = 0.4379, treatment p = 0.0251, diet p = 0.01. Sidak’s post hoc, HFD vs HFD-Succ p = 0.4358, CC vs CC-Succ p = 0.058). **H)** Microglia soma volume (Two-way ANOVA, diet*treatment interaction p = 0.5314, treatment p = 0.0191, diet p = 0.0071. Sidak’s post hoc, HFD vs HFD-Succ p = 0.3284, CC vs CC-Succ p = 0.0586). **I)** Microglia soma sphericity (Two-way ANOVA, diet*treatment interaction p = 0.8121, treatment p = 0.8647, diet p = 0.7008). N= 5-8/group (A-B), N=3-4/group (C-I) Error bars represent SEM. *p ≤ 0.05, **p ≤ 0.01 and ***p ≤ 0.001, ****p ≤ 0.0001.

Based on the above-mentioned results, we investigated microglia cells given their well-established role in neuroinflammation,^53,54^ and since succinate seem to prevent their conversion to a pro-inflammatory state in vitro.^55^ The density of microglial cells in the hippocampus was not affected by either diet or succinate treatment. (Fig 7C). Microglia complexity, filament lengths, branching points and terminal points were not significantly altered (Fig. 7D-F). On the other hand, there was a main effect of diet on soma area and volume showing a decrease in CC compared to HFD (Fig. 7G-H), while succinate showed a tendency to increase both measures (Fig. 7G-H). Sphericity was not influenced by diet or succinate (Fig. 7I).

All together, these data point toward a modulation of inflammation in the hippocampus by succinate administration, which may underscore the improved behavioural performance.

## DISCUSSION

In this study, we investigated the metabolic and behavioral effects of three dietary interventions -CC, CC-IF and HDF-IF - following DIO in mice. Our findings demonstrate that both CC and IF reversed the metabolic perturbations induced by chronic HFD consumption, albeit with distinct molecular and behavioral profiles. Indeed, CC-IF was exclusive in improving exploration and anxiety-like behavior. CC-IF was characterized by a decrease in plasma succinate and its potential shunt to the BAT and liver, which could influence energy expenditure and inflammatory responses.

As expected, mice fed a HFD for 14 weeks developed obesity, evidenced by significantly increased BW and impaired glucose tolerance. Among the dietary interventions, the CC-IF regimen was the most effective in reducing both BW and plasma insulin levels compared to HFD-IF and CC alone. This is in line with a recent study conducted in elderly individuals comparing IF and a healthy diet.^56^ However, all three regimens successfully restored glucose tolerance. CC-IF also had the most pronounced effect in reducing fat depots. In contrast, the reduction in plasma leptin levels following the dietary switch was comparable across HFD-IF, CC, and CC-IF.

From a translational perspective, these findings suggest that substantial reductions in BW or fat mass may not be required to restore glucose tolerance and lower plasma insulin and leptin levels. Rather, a more moderate weight loss or dietary modulation may suffice to improve key metabolic parameters.

Nonetheless, the liver, a major player in nutrient processing and whole-body metabolic homeostasis,^57^ displayed specific transcriptional responses to the three different diet switches in comparison to ad libitum HFD feeding. CC-IF had the most dramatic impact with 1537 differential expressed transcripts, while CC and HFD-IF accounted for 463 and 577 DEG, respectively. CC-IF upregulated gene top annotations were related to transcriptional regulation. This is in line with the hepatic tissue tendency to save energy in a condition of limited food and energy resources. Moreover, the liver of the CC-IF, CC and HFD-IF groups experienced a decrease in the transcription of genes involved in lipid biosynthetic processes, while circadian rhythms came up in the annotations of upregulated transcripts. Alterations in liver clock function in response to diet have been previously observed and described as an adaptation mechanism upon a nutritional challenge.^58–63^ Although we did not examine daily oscillations in gene expression, our findings suggest that the hepatic clock may play a role in sensing nutritional changes following a dietary switch, independently of the diet specific composition.

In CC-IF mice, gene sets were significantly enriched for pathways involved in chromatin remodeling, transcriptional regulation, protein trafficking, and DNA repair, mechanisms typically associated with cellular homeostasis and adaptability.^49^ In contrast, HFD-IF mice showed enrichment in pathways related to extracellular matrix organization, actin filament dynamics, and responses to mechanical stress, features commonly associated with tissue remodeling and pathological states.^64,65^ These HFD-IF transcriptional signatures may reflect a state of subclinical hepatic inflammation, potentially driven by lipotoxicity.^43^ This could be the consequence of prolonged lipid exposure, both from dietary fat during feeding and from adipose tissue mobilization during fasting days. The cyclic overload of lipids may challenge the liver capacity to maintain lipid homeostasis, ultimately leading to stress responses and inflammation. Thus, while IF may have beneficial effects in the context of a balanced diet, it could exacerbate liver related metabolic dysfunctions when applied in a HF intake context. Intriguingly, GO analysis reveals that only CC-IF, not HFD-IF or CC alone, downregulated ubiquitin-proteasome pathway terms in liver and hippocampus-a cascade essential for protein homeostasis and cellular quality control.^66^ This suggests that the combined intervention featured by CC-IF uniquely reduces the cellular demand for proteolytic clearance, likely reflecting improved proteostasis under this diet.

The CC-IF regimen was particularly effective in improving behavioral outcomes. It was the only intervention that consistently ameliorated both exploratory activity and anxiety-like behavior. With the exception of unsupported rearing, mice switched to the CC diet alone did not show significant differences from HFD-fed animals. The HFD-IF group exhibited some improvements in specific parameters within the OF and EPM tests. Previous studies have shown that IF enhances hippocampal plasticity, promoting long-term potentiation and increasing neurogenesis,^67,68^ which may ultimately contribute to improved behavioral performance. However, neither BW reduction nor the acute effects of IF alone fully explain the distinct behavioral profiles observed among the experimental groups. The emotionality score further confirmed the superior efficacy of the CC-IF intervention from a behavioral standpoint. Finally, a recent human study showed that IF led to better outcomes than a healthy diet in specific executive function tasks in older adults,^56^ which may align with our data.

Our metabolites analysis highlighted succinate as a top hit regulated molecule in CC-IF. Intriguingly, succinate was increased in the liver and BAT and decreased in CC-IF plasma. The reduction in circulating succinate could be protective. Elevated plasma or serum succinate concentration has been found in human obesity compared to lean individuals and the increase seems associated with impaired glucose metabolism.^69,70^ Higher circulating succinate levels consistently correlate with a cluster of established cardiometabolic risk factors, such as BMI, higher fasting insulin, dyslipidemia, increased diastolic blood pressure, and systemic inflammation, as indicated by elevated C-reactive protein.^69^ Why CC-IF influences plasma succinate levels is not yet fully understood. However, it may be the most effective regimen for activating thermogenesis, as suggested by the pronounced reduction in BW. CC-IF could promote, through yet unknown mechanisms, the uptake of succinate by BAT,^71^ where it may be used to stimulate UCP1 activity.^51^ On the other hand, the increase in hepatic succinate levels may reflect its role in supporting gluconeogenesis by replenishing TCA cycle intermediates and sustaining the energy demand required for glucose production. CC-IF could be the diet in which a burst in gluconeogenesis is necessary to avoid hypoglycemia. Liver tissue has also been shown to uptake circulating succinate.^72,73^ Interestingly, during exercise and overnight fasting a significant hepato-splanchnic bed uptake of succinate was observed in human subjects,^74^ in line with a likely succinate liver uptake upon CC-IF. Finally, we can not exclude IF mechanisms involving modulation of intestinal^75^ or gut microbiota derived circulating succinate.^76^ Thus, the BAT, and potentially the liver may contribute through a “metabolic vacuum” effect to reduce plasma succinate levels upon CC-IF. This could be a novel mechanism that distinguishes CC-IF from caloric restriction, granting IF superior capability to counteract adiposity, glucose intolerance, and potentially the effect of HFD-induced dysbiosis,^70,77^ enhancing a reduction in cardiometabolic risk.

A pivotal aspect of our study was exploring whether the unique succinate signature present in CC-IF mice could be linked to the observed benefits. To test this, we administered sodium succinate in drinking water to DIO mice. Strikingly, succinate supplementation recapitulated several key beneficial outcomes observed in CC-IF. Metabolically, while HFD-Succ mice showed only a slight, non-significant mitigation of HFD-induced weight gain, CC-Succ mice exhibited a significant reduction in BW compared to the CC group, mirroring the enhanced weight loss seen with CC-IF. This suggests that succinate effect on BW is context-dependent, possibly being more effective when combined with a healthier background diet. Importantly, succinate significantly improved glucose tolerance in HFD-fed mice, aligning with previous studies demonstrating succinate ability to enhance glucose homeostasis by promoting BAT thermogenesis and energy expenditure.^51,78,79^ While Gaspar et al. (2023) also showed succinate protects against DIO, our data contrast slightly with Ives et al. (2020), who reported reduced adiposity without clear glucose tolerance improvement.^80^ The most pronounced improvement in glucose clearance was seen in the CC-Succ group, again paralleling the CC-IF results and suggesting synergy between succinate and a balanced diet. Furthermore, succinate influenced fat depot distribution, particularly BAT weight, consistent with its known role in BAT activation.^51,81^

Behaviorally, succinate supplementation exerted effects remarkably similar to those of CC-IF. Indeed, the emotionality score confirmed that CC-Succ group significantly reduced anxiety-like behavior compared to the HFD condition, achieving a profile similar to CC-IF mice. Given the links between DIO, neuroinflammation, cognitive decline, and mood disorders,^19,82,83^ we investigated whether succinate could modulate inflammatory markers in the hippocampus. Our transcriptomic analysis in HFD, CC-IF, HFD-IF and CC hippocampal tissue revealed that HFD feeding induced pathways related to immune response and neuroinflammation. Notably, succinate supplementation significantly reduced the expression of the pro-inflammatory cytokines IL-1β and TNFα in the hippocampus of CC-Succ mice compared to CC mice, and for TNFα compared to HFD, suggesting a direct anti-inflammatory effect within the brain or an indirect effect mediated by improved peripheral metabolism. This finding is particularly interesting given the often-described pro-inflammatory role of succinate accumulation within activated immune cells like macrophages, where it stabilizes HIF-1α and drives IL-1β production.^84–86^ However, extracellular succinate signaling via its receptor SUCNR1 can have more complex, context-dependent roles, including anti-inflammatory or resolving actions.^77,85,87,88^ Systemic administration, as used here, likely increases circulating succinate, potentially engaging SUCNR1 on various cell types such as neurons, microglia,^55^ or altering peripheral signals that impact the brain. Our observation aligns with the concept that systemic metabolic shifts (like those induced by IF or potentially succinate supplementation) can dampen central inflammation.^83,89^ Analysis of microglia morphology showed that while switching to CC diet reduced soma size/volume compared to HFD, succinate tended to increase these parameters, perhaps reflecting an altered activation or metabolic state rather than simple quiescence, warranting further investigation.

In summary, our study demonstrates that systemic succinate supplementation, mimicking certain outcomes of CC-IF, exerts beneficial effects on behavior and reduces inflammatory markers in the hippocampus of DIO mice. The mechanisms likely involve a combination of factors: improved peripheral metabolic health indirectly benefiting the brain, and potentially direct central actions including modulation of inflammatory pathways. Dissecting the relative contributions of these peripheral versus central, and metabolic versus signaling, effects of succinate remains a key area for future investigation. Finally, our study reveals that CC-IF induces body site–specific modulation of succinate, highlighting a novel mechanism that may underlie its metabolic benefits compared to a balanced ad libitum diet. These findings suggest that IF in individuals with obesity or other metabolic diseases may offer enhanced protection against cardiometabolic risk, and reduce the likelihood of developing neuropsychiatric disorders or cognitive dysfunction.

## Acknowledgments

We thank the members of Tognini’s team for insightful comments and feedback. Special thanks to Giulia Tiozzo for her help with succinate experiments, to Sara and Cecilia Ciampi and Dr. Silvia Burchielli for their help in the mouse facility. This research was supported in part by NextGenerationEU Italian Ministry of University and Research M4.C2 -PNRR YOUNG MSCA_0000081 iNsPIReD, and PRIN-PNRR2022 CARE P2022CXN7X CUP I53D23006920001 to P.T; in part through an EFSD award supported by EFSD/Lilly European Diabetes Research Programme to AG and PT, and in part through the European Union’s Horizon Europe Research and Innovation Programme for the project “PAS GRAS: under grant agreement No. 101080329 to A.G.

## Author contributions

A.T. performed all the experiments, analyzed the data, and prepared the figures. F.C., performed all the metabolome analysis and analysed the data. S.A. and K.C.A. performed the RNA-seq analysis. S.C., M.G.G., A.M., F.D. and M.N. helped with tissue collection, behavioural and molecular/immunofluorescence experiments. L.dB. quantified succinate in BAT G.S. and M.M. collected adipose tissues. P.B. supervised the RNA-seq analysis and interpreted the data. A.G. supervised, analysed and interpreted the metabolomics. M.M., A.G. and P.B. gave feedback on the manuscript. P.T. and A.G. conceived the project. P.T. designed, supervised and performed the experiments, interpreted the data, and wrote the manuscript.

## MATERIALS AND METHODS

### Animals and Feeding

All the experiments were carried out in accordance with the European Directives (2010/63/EU) and were approved by the Italian Ministry of Health.

7-8 week-old male C57BL/6J mice (JAX #00064) were housed (2-3 per cage) in individually ventilated cages in a 12-hour light/12-hour dark environment at a controlled temperature (23°C). All the mice were initially fed ad libitum a high fat diet (60% Kcal from fat, irradiated Research Diet #D12492) for 10 weeks and then divided in 4 different feeding groups for the next 4 weeks: a group continuing on the HFD (HFD group), a group practicing IF by consuming the HFD every other day (HFD-IF), a group fed a balanced control chow diet ad libitum (CC, irradiated Research Diet #12450J), and a group practicing IF by consuming the control chow every other day (CC-IF).

At the end of the 14 weeks of dietary regimen, 12 mice/experimental group were sacrificed by decapitation. Half of the animals were used for the RNA extraction experiment, leptin, and insulin, OGTT analysis. Half of the animals were used for metabolomics analysis. Hippocampus, liver, subcutaneous fat, perirenal fat, interscapular brown adipose tissue (BAT) and epididymal fat were harvested, snapped frozen in liquid nitrogen and stored at - 80°C until use. Fresh blood was centrifuged at 1500×g for 15 minutes and plasma isolated and stored at -80°C until use.

### Oral glucose tolerance test (OGTT)

The OGTT was performed at the end of the 13th week of diet treatments. Before the test, mice were housed in clean cages and fasted for 6 hours. Mice were given an oral bolus of glucose (2 g/kg of body weight; D-glucose, Sigma Aldrich Catalog #G6152) and glycemia from tail vein blood samples was quantified through a glucometer (Accu Chek Guide). Glycemia was evaluated at 0, 15, 30, 60, 120 minutes post glucose administration.

### Plasma insulin and leptin levels

Plasma insulin and leptin levels were analysed using the mouse metabolic hormone panel (Merck Catalog #MMHE-44K), following manufacturer instructions.

### Behavioral tests

Behavioral performance was assessed in a dedicated cohort of mice (n = 8/11 experimental group). During the last 7 days of diet feeding, mice underwent the open field (OF) test, the novel object recognition test (NORT), the Y maze test and the elevated plus maze test (EPM).

#### Open Field (OF) test and Novel Object Recognition Test (NORT)

In our battery of tests, the OF has been exploited for the NORT habituation phase.

The apparatus consisted of a square arena (60 × 60 × 30 cm) constructed in poly(vinyl chloride) with black walls and a white floor. The mice received one session of 10-min duration in the empty arena to habituate them to the apparatus and test room. Animal position was continuously recorded by a video tracking system (Noldus Ethovision XT). In the recording software an area corresponding to the center of the arena (a central square 30 × 30 cm), and a peripheral region (corresponding to the remaining portion of the arena) were defined. The total movement of the animal and the time spent in the center or in the periphery area were automatically computed. Mouse activity during this habituation session was analyzed for evaluating the behavior in the OF arena. Rearing behavior, measured as the number of times in which an animal temporarily stands on its hindlimbs, was manually counted and used as an exploratory and anxiety-like behavior index.^90^

The NORT consisted of two phases: sample and testing phase. During the sample phase, two identical objects were placed in diagonally opposite corners of the arena, approximately 6 cm from the walls, and mice were allowed to explore the objects for 10 minutes, then they were returned to their cage. The objects to be discriminated against were made of glass and were too heavy to be displaced by the mice. The testing phase was performed 24 h after the sample phase. One of the two familiar objects was replaced with a new one and the mice were allowed to explore objects for 5 min. To avoid possible preferences for one of two objects, the choice of the new and old object and the position of the new one were randomized among animals. The amount of time spent exploring each object (nose sniffing and head orientation within <2.0 cm) was recorded and evaluated by the Noldus Ethovision XT video tracker system. Arena and objects were cleaned with 70% ethanol between trials to stop the build-up of olfactory cues. A preference index was computed as PI (%) = (T new/(T new + T familiar)x100, where T new is the time spent exploring the new object, and T familiar is the time spent exploring the old one. Any mouse that explored for less than 13 seconds was excluded from the analysis.

#### Y maze test

A Y-shaped maze with three symmetrical gray solid plastic arms at a 120-degree angle (26 cm length, 10 cm width, and 15 cm height) was used to test exploratory and hyperactive behaviors. Mice were placed in the center of the maze and allowed to freely explore the maze for 8 minutes. The apparatus was carefully cleaned with ethanol between trials to avoid the build-up of odor traces. All sessions were video-recorded (Noldus Ethovision XT) for offline blind analysis. The arm entry is defined as all four limbs within the arm. A triad is defined as a set of three arm entries, when each entry is to a different arm of the maze. The number of arm entries and alternate triads were recorded in order to calculate the alternation percentage (generated by dividing the number of alternate triads by the number of total triads and then multiplying by 100), as explained in.^91^

#### Elevated plus maze (EPM) test

The plus shaped arena was installed on a raised support and a camera (Noldus Ethovision XT) was used to monitor the mice movements. The arena consisted of a total of four arms: two open and two enclosed by Plexiglas walls (50 cm long and 10 cm wide) with the same illumination levels. Mice were placed in the center of the EPM arena and exploration was video-tracked for 5 min. Time spent in open and closed arms and frequency of entries were analyzed, as well as the mean velocity and the total distance traveled by each animal. The percentage of time spent in the open arm was evaluated as (T open/(T open + T closed)x100, where T open is the time spent in the open arm, and T closed is the time spent in the closed arm. The percentage of entries in the open arm was computed as (N° open/(N° open + N° closed)x100, where N° open is the number of entries in the open arm, and N° closed is the number of entries in the closed arm. The EPM arena was cleaned with 70% ethanol between each trial. Mice that fell from the platform were excluded from the analysis.

The emotionality score was calculated as explained in.^47^ It was derived by averaging the z-scores of four parameters: time spent in the center (#1) and the number of total rears (#2) in the OF test, along with percentage of time spent (#3) and the percentage of number of entries (#4) in the open arm during the EPM. Animals that were excluded from the behavioural tests used for calculation of the emotionality score were also excluded from the score.

### RNA extraction

Hippocampus and liver samples were homogenized in phenol/guanidine-based QIAzol Lysis Reagent (Qiagen #79,306). Chloroform was added and the samples were shaken for 15 s.

The samples were left at 20–24 °C for 3 min and then centrifuged (12,000×g, 20 min, 4 °C). The upper phase aqueous solution, containing RNA, was collected in a fresh tube and the RNA was precipitated by the addition of isopropanol. Samples were mixed by vortexing, left at RT for 10 min and then centrifuged (12,000×g, 15 min, 4 °C). The supernatant was discarded and the RNA pellet was washed in 75% ethanol by centrifugation (7500×g, 5 min, 4 °C). The supernatant was discarded and the pellet was left to dry for a minimum of 15 min before resuspending in RNAse free water.

RNA concentration was determined by the Nanodrop Spectrophotometer (Thermoscientific 2000 C). RNA quality was analyzed via agarose gel electrophoresis (1% agarose).

### qPCR

Total RNA was reverse transcribed using QuantiTech Reverse Transcription Kit (Qiagen Cat. # 205311). Gene expression was analyzed by real-time PCR (Step one, Applied Biosystems), using PowerUp SYBR Green Master Mix (Thermo Fisher #A25742). The primers for gene expression were designed by Primer 3 software (v. 0.4.0), and are reported below:

*Actb:* Forward: 5’-GGCTGTATTCCCCTCCATGC-3’;

*Actb:* Reverse: 5’-CCAGTTGGTAACAATGCCATGT-3’;

*Tnfα:* Forward: 5’-CCCATATACCTGGGAGGAGTCTTC-3’;

*Tnfα:* Reverse: 5’-CATTCCCTTCACAGAGCAATGAC-3’;

*Il1ꞵ:* Forward: 5’-AGTTGACGGACCCCAAAAG-3’;

*Il1ꞵ:* Reverse: 5’-AGCTGGATGCTCTCATCAGG-3’.

Quantitative values for cDNA amplification were calculated from the threshold cycle number (Ct) obtained during the exponential growth of the PCR products. Threshold was set automatically by the Step one software. Data were analyzed by the ΔΔCt methods using the *actb* expression to normalize the cDNA levels of the transcripts under investigation.

### RNA sequencing (seq)

Library preparation and RNA-seq technique were performed by a commercially available service (IGATech Udine, Italy). Total RNA was extracted as described above, libraries were prepared and sequenced on Illumina HiSeq2500 instrument during a pair-end read 125 bp sequencing, producing sequencing results in FastQ format. The FastQ files were processed through the standard Tuxedo protocol, using Tophat and Cufflinks. Tophat was used to align the RNA-seq reads to the reference genome assembly mm10 and Cufflinks was used to calculate gene expression levels. This protocol outputs the FPKM values for each gene of each replicate.

The differential analysis of the FPKM values across all experiments and control groups was conducted with Cyber-T, a differential analysis program using a Bayesian-regularized t-test.^92,93^ The p-value thresholds used to determine differential expression were 0.05 for the liver and 0.1 for the hippocampus.

### Gene ontology analysis

Gene ontology analysis was performed using Database for Annotation, Visualization and Integrated Discovery (DAVID) v6.7, using genomic background; GO Biological Processes were chosen for gene clustering.

### Metabolomics

Polar metabolites were quantified in 40 µl of plasma and in about 20 mg of liver, hippocampus and brown adipose tissue (BAT). Tissues were homogenized using Precellys Evolution Homogenizer (Bertin Instruments, Frankfurt, Germany). Samples were deproteinized with methanol by centrifugation for 20 minutes at 13300 rpm at 4°C and metabolites were extracted by a modified Folch method^94^ using chloroform:methanol (2:1) and water, after 15 minutes of centrifugation at 13300 rpm at 4°C. Upper phases were dried under gentle nitrogen flux and were derivatized using 10 µl of methoxyamine hydrochloride (Merck KGaA, Darmstadt, Germania) solution in pyridine (20 mg/mL) for 30 minutes at 60°C and 30 µl of N-(tert-butyldimethylsilyl-N-methyltrifluoroacetamide (TBDMS, Merck KGaA, Darmstadt, Germania) with 70 µl acetonitrile for 1 hour at 60°C. Amino acids and organic acids were quantified using labeled internal standard mix (MSK-A2-S Metabolomics Amino Acid Mix standard, CIL Cambridge, MA, USA and MSK-OA-1 Labeled Organic Acid Mix, CIL Cambridge, MA, USA), and measured by gas chromatography/tandem mass spectrometry (GC 8890/MS 7000D, Agilent, Santa Clara, CA).

### Succinate treatment

A dedicated cohort of 7-8 week-old male C57BL/6J mice were initially fed ad libitum a HFD for 10 weeks and then divided in 4 different groups for the next 4 weeks: a group continuing on the HFD (HFD), a group consuming HFD and drinking Succinate (HFD-Succ), a group fed a CC diet ad libitum (CC), and a group consuming CC and drinking Succinate (CC-Succ). Sodium succinate (Sigma Aldrich #224731) was dissolved in drinking water (1.5% w/vol) and refreshed every 2 days. A total of 11-14 mice/experimental groups were used in this experiment. Half of the animals were sacrificed and fresh tissues collected. Half of the animals were transcardially perfused.

### Microglia immunofluorescence analysis

Mice were anesthetized with chloral hydrate (20ml/Kg BW) and perfused via intracardiac infusion with PBS and then 4% paraformaldehyde (PFA, w/vol, dissolved in 0.1 M phosphate buffer, pH 7.4). Brains were quickly removed and post-fixed overnight in PFA at 4 °C, then transferred to 30% sucrose (w/vol) solution. 45 μm coronal sections were cut on a freezing microtome (Leica) and free-floating sections were processed for immunofluorescence.

Hippocampal sections were incubated for 2 h in a blocking solution containing 10% BSA (w/vol) and 0.5% Triton X-100 (vol/vol) in PBS, and incubated overnight at 4 °C with guinea pig anti-iba1 (Synaptic Systems # 234308) diluted 1:500 in PBS with 3% BSA (w/vol) and 0.15% Triton X-100 (vol/vol).

Sections were then washed with PBS and incubated for 2h at 22-24 °C with Alexa Fluor 488–conjugated secondary antibody (Invitrogen # A-11073), which was added at a dilution of 1:500 in the same solution as the primary antibody.

Sections were washed three times with PBS and mounted on slides, then they were air-dried and coverslipped with Vectashield mounting medium (Vector Laboratories, Cat. no. H-1000). Imaging was performed on an Leica STELLARIS 5 confocal microscope (Leica Microsystem, Wetzlar, Germany) using a dry 20x and a 63x oil objective. The area of the hippocampus was defined based on the mouse brain atlas (Paxinos and Franklin’s the Mouse Brain in Stereotaxic Coordinates). IBA1 positive cell counting for microglia density analysis was performed using a custom MATLAB-based graphical user interface.^95^

The 3D reconstruction of microglial cells was performed on Z-stacks of ∼ 45 µm, acquired with a z-step of 0.50 μm (for a final voxel size of 0.12025 × 0.12025 × 0.5 µm). Images were then processed for filament and soma reconstruction using respectively the “Filaments” and “Surfaces” functions of IMARIS software (Bitplane). The analyzed morphological features were selected based on their relevance as indicators of microglia activation: filament length, number of branching points, number of terminal points, soma area, soma volume and soma sphericity (roundness).^91,96^

Between 4 and 7 microglia cells per mouse were reconstructed for the arborization analysis. Between 7 and 11 cells were used for the soma analysis.

### Statistical analysis

The sample sizes (n) were based on prior studies and are indicated in the figure legend for each panel. Whenever possible, quantification and analyses were performed blind to the experimental condition. The reported “n” refers to individual animals, and each circle over the graph bars represents a single mouse, unless otherwise stated.

Most statistical analyses were performed using GraphPad Prism version 7 (GraphPad Software, San Diego, CA, USA). Outliers were identified using the Grubbs’ test for single outliers, with a significance level set at α = 0.05. The intergroup difference was determined by Two-way analysis of variance (ANOVA), followed by post hoc t-test for multiple comparisons, as specified in each figure legend. Only the P values of statistically significant post-hoc comparisons are reported in the figure legend. All data are represented as the mean ± SEM. P values of ≤0.05 were considered as statistically significant, and significance was defined in the figure panels as follows: *p ≤ 0.05, **p ≤ 0.01 and ***p ≤ 0.001, ****p ≤ 0.0001.

