## Supplementary Figures for "Succinate Modulation as a Novel Mechanism Underlying the Effects of Intermittent Fasting on Brain Function and Metabolism in Diet-Induced Obesity"

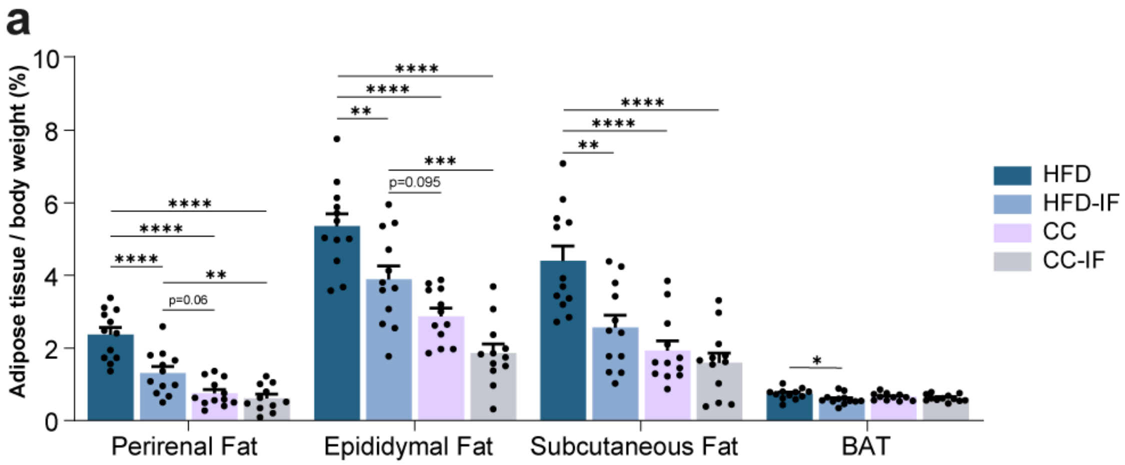


**Suppl. Figure 1:** Perirenal adipose tissue percentage (Two-way ANOVA, diet*feeding mode, interaction $p = 0.0046$, feeding mode $p = 0.0004$, diet $p < 0.0001$. Tukey's post hoc, HFD vs CC-IF $p < 0.0001$, HFD vs CC $p < 0.0001$, HFD vs HFD-IF $p < 0.0001$, CC-IF vs HFD-IF $p = 0.0154$, CC vs HFD-IF $p = 0.0619$). Epididymal adipose tissue percentage (Two-way ANOVA, diet*feeding mode, interaction $p = 0.4694$, feeding mode $p = 0.0002$, diet $p < 0.0001$. Tukey's post hoc, HFD vs CC-IF $p < 0.0001$, HFD vs CC $p < 0.0001$, HFD vs HFD-IF $p = 0.0073$, CC-IF vs HFD-IF $p = 0.0001$, CC vs HFD-IF $p = 0.1012$, CC vs CC-IF $p = 0.0952$). Subcutaneous adipose tissue percentage (Two-way ANOVA, diet*feeding mode, interaction $p = 0.0281$, feeding mode $p = 0.002$, diet $p < 0.0001$. Tukey's post hoc, HFD vs CC-IF $p < 0.0001$, HFD vs CC $p < 0.0001$, HFD vs HFD-IF $p = 0.0016$). Brown adipose tissue percentage (Two-way ANOVA, diet*feeding mode, interaction $p = 0.1236$, feeding mode $p = 0.0244$, diet $p = 0.5088$. Tukey's post hoc, HFD vs HFD-IF $p = 0.0376$). N = 11-12/experimental group. Error bars represent SEM. *$p < 0.05$, **$p < 0.01$, ***$p < 0.001$, ****$p < 0.0001$.

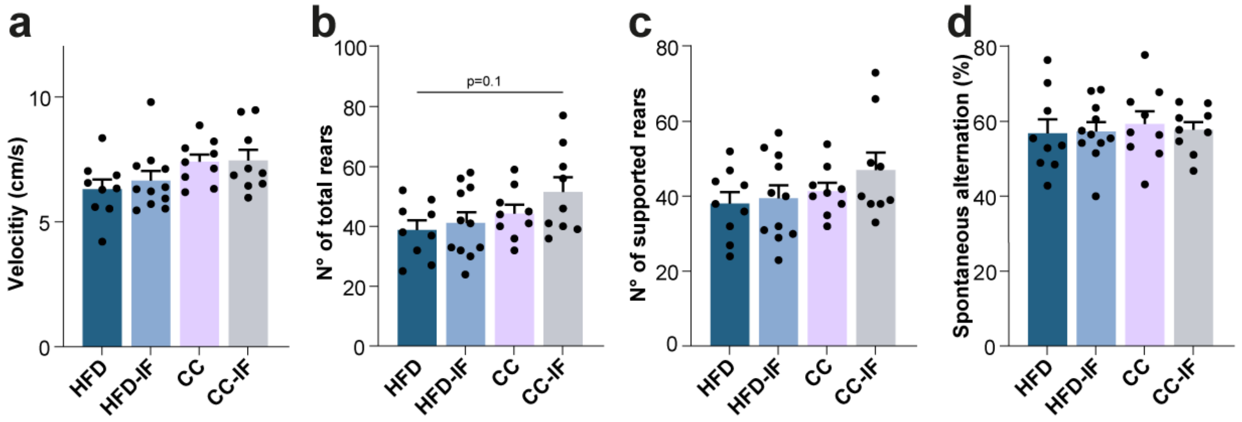


**Suppl. Figure 2: A)** Velocity in the Open Field test (Two-way ANOVA, diet*feeding mode, interaction $p = 0.7012$, feeding mode $p = 0.6033$, diet $p = 0.0173$). **B)** Total number of rears in the Open Field test (Two-way ANOVA, diet*feeding mode, interaction $p = 0.5276$, feeding mode $p = 0.2057$, diet $p = 0.0392$. Tukey's post hoc, HFD vs CC-IF $p = 0.1018$). **C)** Number of supported rears in the Open Field test (Two-way ANOVA, diet*feeding mode, interaction $p = 0.5456$, feeding mode $p = 0.313$, diet $p = 0.1252$). **D)** Spontaneous alternation percentage in the Y-maze test (Two-way ANOVA, diet*feeding mode, interaction $p = 0.7364$, feeding mode $p = 0.8494$, diet $p = 0.6229$. Tukey's post hoc, HFD vs CC-IF $p = 0.0707$, CC vs CC-IF $p = 0.0654$). N = 9-11/experimental group. Error bars represent SEM. *$p < 0.05$, **$p < 0.01$, ***$p < 0.001$, ****$p < 0.0001$.

**a**

| Comparison | up in HFD | down in HFD |
|---|---|---|
| HFD vs CC-IF | 90 | 28 |
| HFD vs CC | 60 | 13 |
| HFD vs HFD-IF | 114 | 21 |

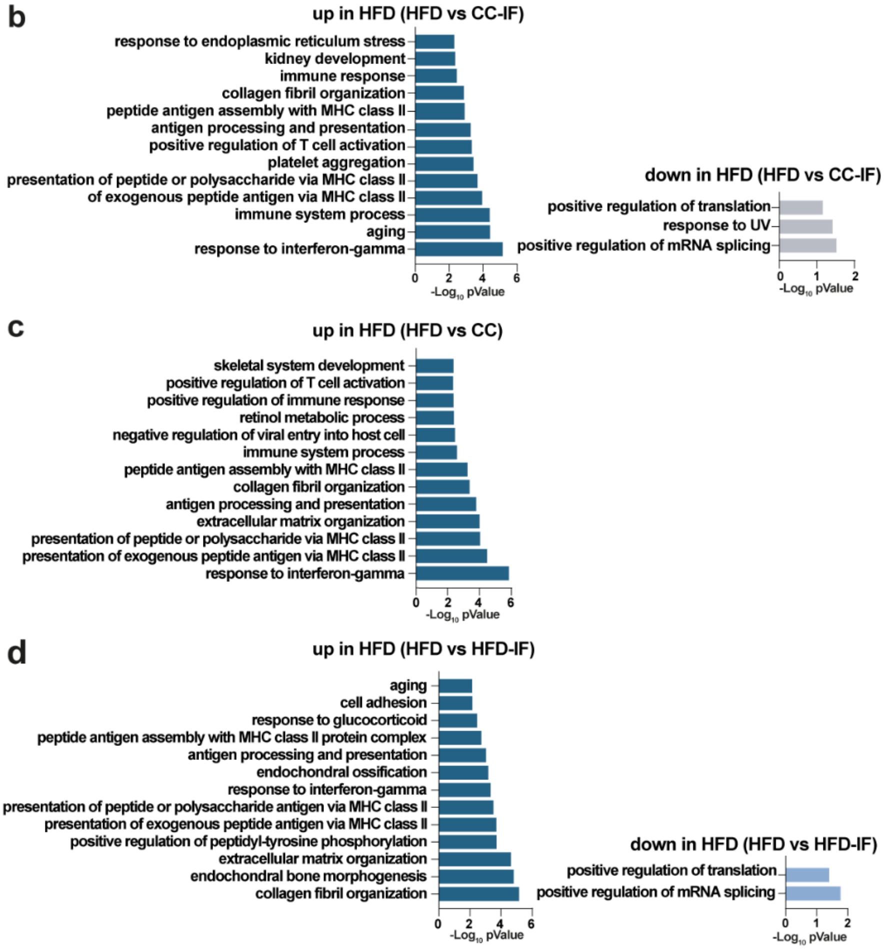


**Suppl. Figure 3: A)** Summary table illustrates the number of DEGs (pvalue < 0.1) between each of the 3 dietary interventions (HFD-IF, CC, CC-IF) in comparison with the HFD in the hippocampus. **B)** Gene ontology analysis (biological processes) of 90 DEGs upregulated in HFD vs CC-IF, top 13 significant pathways, pValue ranked (left). Gene ontology analysis biological processes of 28 DEGs upregulated in CC-IF vs HFD, pValue ranked (right). **C)** Gene ontology analysis (biological processes) of 60 DEGs upregulated in HFD vs CC, top 13 significant pathways, pValue ranked. **D)** Gene ontology analysis (biological processes) of 114 DEGs upregulated in HFD vs HFD-IF, top 13 significant pathways, pValue ranked (left). Gene ontology analysis (biological processes) of 21 DEGs upregulated in HFD-IF vs HFD, pValue ranked (right).

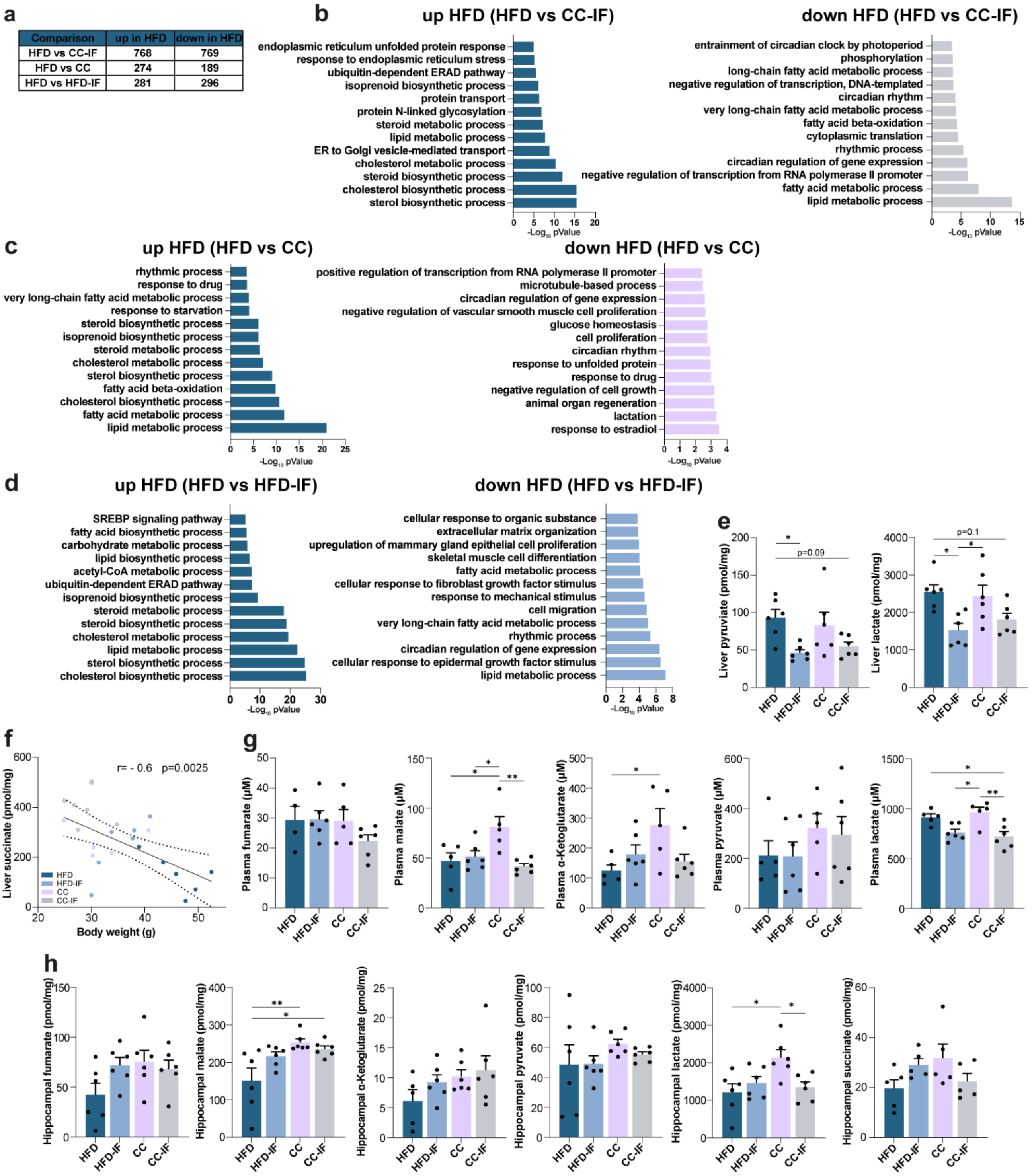


**Suppl. Figure 4: A)** Summary table illustrates the number of DEGs (pValue < 0.05) between each of the 3 dietary interventions (HFD-IF, CC, CC-IF) in comparison with the HFD in the liver. **B)** Gene ontology analysis biological processes of 768 DEGs upregulated in HFD vs CC-IF, top 13 significant pathways, pValue ranked (left). Gene ontology analysis biological processes of 769 DEGs upregulated in CC-IF vs HFD, top 13 significant pathways, pValue ranked (right). **C)** Gene ontology analysis biological processes of 274 DEGs upregulated in HFD vs CC, top 13 significant pathways, pValue ranked (left). Gene ontology analysis biological processes of 189 DEGs upregulated in CC vs HFD, top 13 significant pathways, pValue ranked (right). **D)** Gene ontology analysis biological processes of 281 DEGs upregulated in HFD vs HFD-IF, top 13 significant pathways, pValue ranked (left). Gene ontology analysis biological processes of 296 DEGs upregulated in HFD-IF vs HFD, top 13 significant pathways, pValue ranked (right). **E)** Pyruvate concentration in liver (Two-way ANOVA, diet*feeding mode, interaction p = 0.3989, feeding mode p = 0.0026, diet p = 0.9437. Tukey's post hoc, HFD vs CC-IF p = 0.0934, HFD vs HFD-IF p = 0.0301). Lactate concentration in liver (Two-way ANOVA, diet*feeding mode, interaction p = 0.3686, feeding mode p = 0.001, diet p = 0.7133. Tukey's post hoc, HFD vs CC-IF p = 0.098, HFD vs HFD-IF p = 0.0148, CC vs HFD-IF p = 0.034). **E)** Correlation analysis between Succinate concentration in plasma and body weight (n = 23. Pearson r = - 0.6, p = 0.0025). **G)** Fumarate concentration in plasma (Two-way ANOVA, diet*feeding mode, interaction p = 0.2918, feeding mode p = 0.336, diet p = 0.2533). Malate concentration in plasma (Two-way ANOVA, diet*feeding mode, interaction p = 0.0063, feeding mode p = 0.0241, diet p = 0.1086. Tukey's post hoc, HFD vs CC p = 0.0217, CC vs HFD-IF p = 0.0402, CC vs CC-IF p = 0.005). α-Ketoglutarate concentration in plasma (Two-way ANOVA, diet*feeding mode, interaction p = 0.0189, feeding mode p = 0.3475, diet p = 0.0771. Tukey's post hoc, HFD vs CC p = 0.0342, CC vs CC-IF p = 0.0929). Pyruvate concentration in plasma (Two-way ANOVA, diet*feeding mode, interaction p = 0.8455, feeding mode p = 0.8173, diet p = 0.1412. Lactate concentration in plasma (Two-way ANOVA, diet*feeding mode, interaction p = 0.3306, feeding mode p = 0.0002, diet p = 0.9238. Tukey's post hoc, HFD vs CC-IF p = 0.0237, CC vs HFD-IF p = 0.0178, CC vs CC-IF p = 0.0046). **H)** Fumarate concentration in the hippocampus (Two-way ANOVA, diet*feeding mode, interaction p = 0.0853, feeding mode p = 0.2811, diet p = 0.1645). Malate concentration in the hippocampus (Two-way ANOVA, diet*feeding mode, interaction p = 0.0414, feeding mode p = 0.2237, diet p = 0.0047. Tukey's post hoc, HFD vs CC-IF p = 0.0245, HFD vs CC p = 0.0058). α-Ketoglutarate concentration in the hippocampus (Two-way ANOVA, diet*feeding mode, interaction p = 0.5429, feeding mode p = 0.2468, diet p = 0.096). Pyruvate concentration in the hippocampus (Two-way ANOVA, diet*feeding mode, interaction p = 0.6192, feeding mode p = 0.6576, diet p = 0.192). Lactate concentration in the hippocampus (Two-way ANOVA, diet*feeding mode, interaction p = 0.0134, feeding mode p = 0.1825, diet p = 0.0455. Tukey's post hoc, HFD vs CC p = 0.0131, CC vs CC-IF p = 0.0411). Succinate concentration in the hippocampus (Two-way ANOVA, diet*feeding mode, interaction p = 0.0291, feeding mode p = 0.9809, diet p = 0.4841). N = 4-6/experimental group. Error bars represent SEM. *p < 0.05, **p < 0.01, ***p < 0.001, ****p < 0.0001.

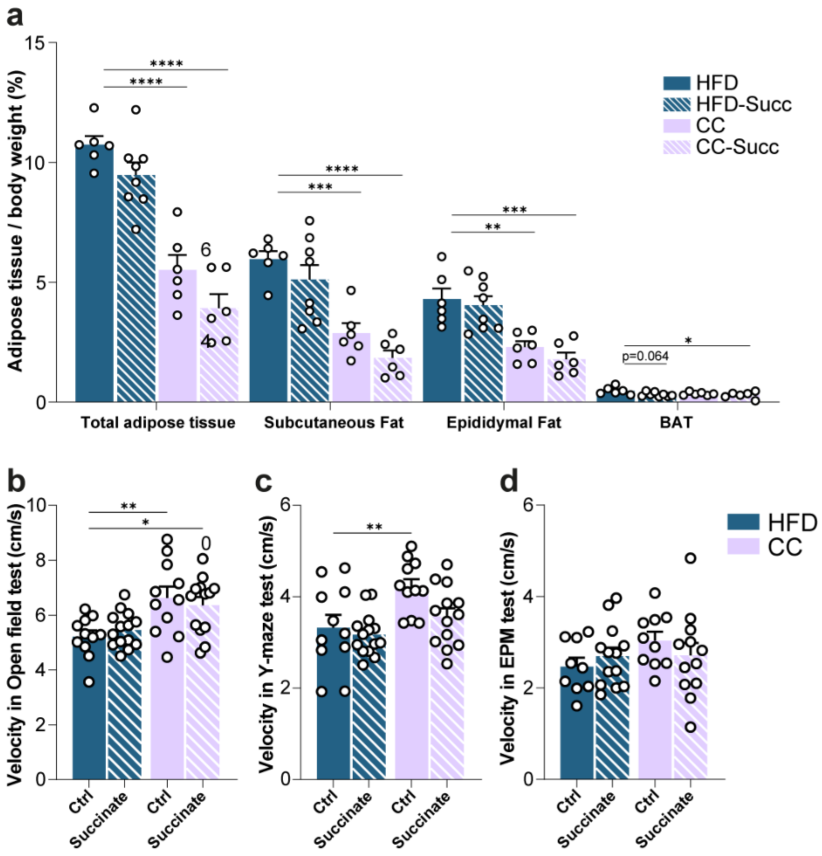


**Suppl. Figure 5: A)** Total adipose tissue percentage (Two-way ANOVA, diet*treatment, interaction p = 0.7653, treatment p = 0.0148, diet p < 0.0001. Dunnett's post hoc, HFD vs CC p < 0.0001, HFD vs CC-Succ p < 0.0001). Subcutaneous adipose tissue percentage (Two-way ANOVA, diet*treatment, interaction p = 0.8579, treatment p = 0.0605, diet p < 0.0001. Dunnett's post hoc, HFD vs CC p < 0.0006, HFD vs CC-Succ p < 0.0001). Epididymal adipose tissue percentage (Two-way ANOVA, diet*treatment, interaction p = 0.7278, treatment p = 0.3076, diet p < 0.0001. Dunnett's post hoc, HFD vs CC p = 0.0025, HFD vs CC-Succ p = 0.0003). Brown adipose tissue percentage (Two-way ANOVA, diet*treatment, interaction p = 0.2968, treatment p = 0.031, diet p = 0.1133. Dunnett's post hoc, HFD vs CC p = 0.064, HFD vs CC-Succ p = 0.0381). **B)** Time spent in the center in the OF (Two-way ANOVA, diet*treatment, interaction p = 0.3655, treatment p = 0.9774, diet p = 0.0001. Dunnett's post hoc, HFD vs HFD-Succ p = 0.8581, HFD vs CC p < 0.0036, HFD vs CC-Succ p = 0.0149). **C)** Velocity in the Y-maze (Two-way ANOVA, diet*treatment, interaction p = 0.214, treatment p = 0.0448, diet p = 0.0018. Dunnett's post hoc, HFD vs HFD-Succ p = 0.8884, HFD vs CC p = 0.0094, HFD vs CC-Succ p = 0.7015). **D)** Velocity in the EPM test (Two-way ANOVA, diet*treatment, interaction p = 0.2158, treatment p = 0.8256, diet p = 0.1982). (A) n = 6-8/experimental group, (B-D) n = 11-14/experimental group. Error bars represent SEM. *p < 0.05, **p < 0.01, ***p < 0.001, ****p < 0.0001.
